# Tumor-induced species-specific dysbiosis drives renal innate immunity and nephrogenic ascites

**DOI:** 10.64898/2026.03.10.710910

**Authors:** Anindita Barua, Fei Cong, Hongcun Bao, Wu-Min Deng

## Abstract

Ascites is a life-threatening complication of advanced malignancies, yet how tumors disrupt systemic fluid homeostasis remains poorly understood. Here, using a *Drosophila* tumor allograft model that recapitulates key features of cancer-associated ascites, we identify a tumor–microbiome–renal axis that controls host fluid balance. Tumor-bearing hosts develop severe abdominal fluid accumulation accompanied by marked expansion and systemic dissemination of the gut commensal *Acetobacter aceti*. Tumor-induced bacterial dissemination activates innate immune signaling in the Malpighian tubules, the insect renal tubules, leading to uric acid accumulation, nephrolithiasis, and progressive ascites. Selective elimination of *A. aceti*, or renal-tubule-specific suppression of IMD/NF-κB signaling, abolishes these pathological changes. Conversely, mono-association of axenic hosts with *A. aceti* is sufficient to recapitulate the ascites phenotype through IMD pathway activation. Together, these findings demonstrate that tumors can remotely induce nephrogenic pathology through species-specific microbiome-dependent immune activation, establishing a mechanistic link between cancer progression and systemic fluid imbalance.

## Introduction

Ascites is characterized by pathological fluid accumulation within the peritoneal cavity, the space between the parietal and visceral peritoneal layers, and can arise from multiple etiologies, including liver cirrhosis, pancreatitis, peritoneal infection, and nephrotic syndrome (1). In addition to these conditions, ascites frequently occurs in cancer patients, particularly in gastrointestinal malignancies (2) and ovarian cancer, where it is most prominent in advanced disease but is also present in approximately one-third of patients at diagnosis (3). Although malignant ascites is clinically common and strongly associated with poor prognosis, the systemic mechanisms by which tumors remotely disrupt host fluid homeostasis remain poorly understood.

Patients with advanced malignancies also frequently develop renal dysfunction. While renal injury in cancer is often attributed to urinary tract obstruction, tumor compression, or renal infiltration (4, 5), a subset of renal pathologies arises independently of direct tumor burden and instead reflects systemic tumor–host signaling. These paraneoplastic renal syndromes are driven by circulating immune mediators, cytokines, and growth factors that alter kidney function (6, 7). Renal dysfunction is strongly associated with fluid imbalance and can contribute to ascites formation; nephrogenic ascites has been reported in chronic kidney disease patients (8, 9). Together, these observations raise the possibility that tumor-induced renal dysfunction contributes directly to cancer-associated ascites. However, the *in vivo* mechanisms connecting tumor growth, host immunity, the microbiome, and renal dysfunction remain difficult to dissect, in part due to limited integrative models.

*Drosophila* tumor models have been instrumental in uncovering systemic consequences of tumor growth, including cachexia-like tissue wasting, characterized by reduced triglyceride levels and elevated carbohydrate content, partly driven by tumor-derived factors such as Unpaired (Upd), Pvf1, and ImpL2 (10–12). Tumors can also disrupt Malpighian tubule (MT) physiology, the functional equivalent of the renal tubules in flies. For instance, the *esg^ts^>yki^3SA^* gut tumors lead to kidney stone formation, uric acid accumulation (13), and fluid retention through tumor-induced antidiuretic signaling mediated by the hormone ITP isoform F (14). Other fly tumors, like oncogenic *Ras^V12^ and aPKC^ΔN^* overexpressing ovarian cancer and allografted tumors, can cause inflammatory signaling-dependent renal stem cell hyperproliferation (15). Collectively, these studies establish *Drosophila* as a powerful system for dissecting tumor - driven systemic dysfunction and fluid homeostasis.

Emerging work has shown that tumor-associated barrier dysfunction can promote bacterial translocation and systemic immune activation (16); however, the identity of the responsible microbes and their functional contribution to renal and fluid phenotypes remain unclear. We previously developed a tumor allograft model that recapitulates tumor-associated dysfunction in nephrocytes (the fly cells functionally analogous to the mammalian glomerulus) (16). This model originates from the transition zone (TZ) of the larval salivary gland, a region characterized by susceptibility to neoplastic transformation. Persistent activation of Notch signaling in this region drives tumor formation (17), and serial transplantation generates stable high-passage TZ tumors (HPTs) that can be propagated across host generations. Using this platform, we observed that HPT-bearing hosts develop renal defects, including uric acid accumulation and kidney stone formation in MTs. Here we test the hypothesis that species-specific gut dysbiosis functions as an upstream driver of tumor-associated nephrogenic ascites. We find that tumor progression is accompanied by selective expansion of the gut commensal bacterium *Acetobacter aceti*, which promotes systemic immune activation and antimicrobial peptide expression in MTs. Targeted depletion of *A. aceti* suppresses immune activation, reduces uric acid accumulation, and alleviates ascites, while renal-specific inhibition of IMD signaling similarly improves fluid balance. Together, our results define a species-resolved tumor–microbiome–immune–renal axis that mechanistically links gut dysbiosis to nephrogenic fluid imbalance during cancer progression.

## Results

### Tumor induced IMD-dependent renal immune activation and metabolic pathology

High-passage tumor (HPT) hosts were generated by injecting primary salivary gland imaginal ring transition zone tumors (*act^ts^>NICD*) (G0) into the abdomens of female adult flies and serially passaging the resulting tumors into new hosts to obtain stable advanced tumors (Fig 1A). The *act^ts^>NICD* primary tumor comprises a posterior neoplastic region along with hyperplastic tissue in the anterior of the *Drosophila* salivary gland transition zone (17). As tumors progressed, hosts showed markedly reduced lifespan relative to Schneider’s (SD) medium-injected controls (16), and developed conspicuous abdominal distension within ∼7–9 days, consistent with ascites formation (Fig. 1B). We confirmed that abdominal distension reflected fluid accumulation by measuring wet and dry weights. Tumor-bearing flies exhibited a significant increase in water mass, whereas dry mass showed a modest, non-significant reduction, resulting in an elevated water-to-dry ratio (Fig. 1C).

**Fig. 1:**
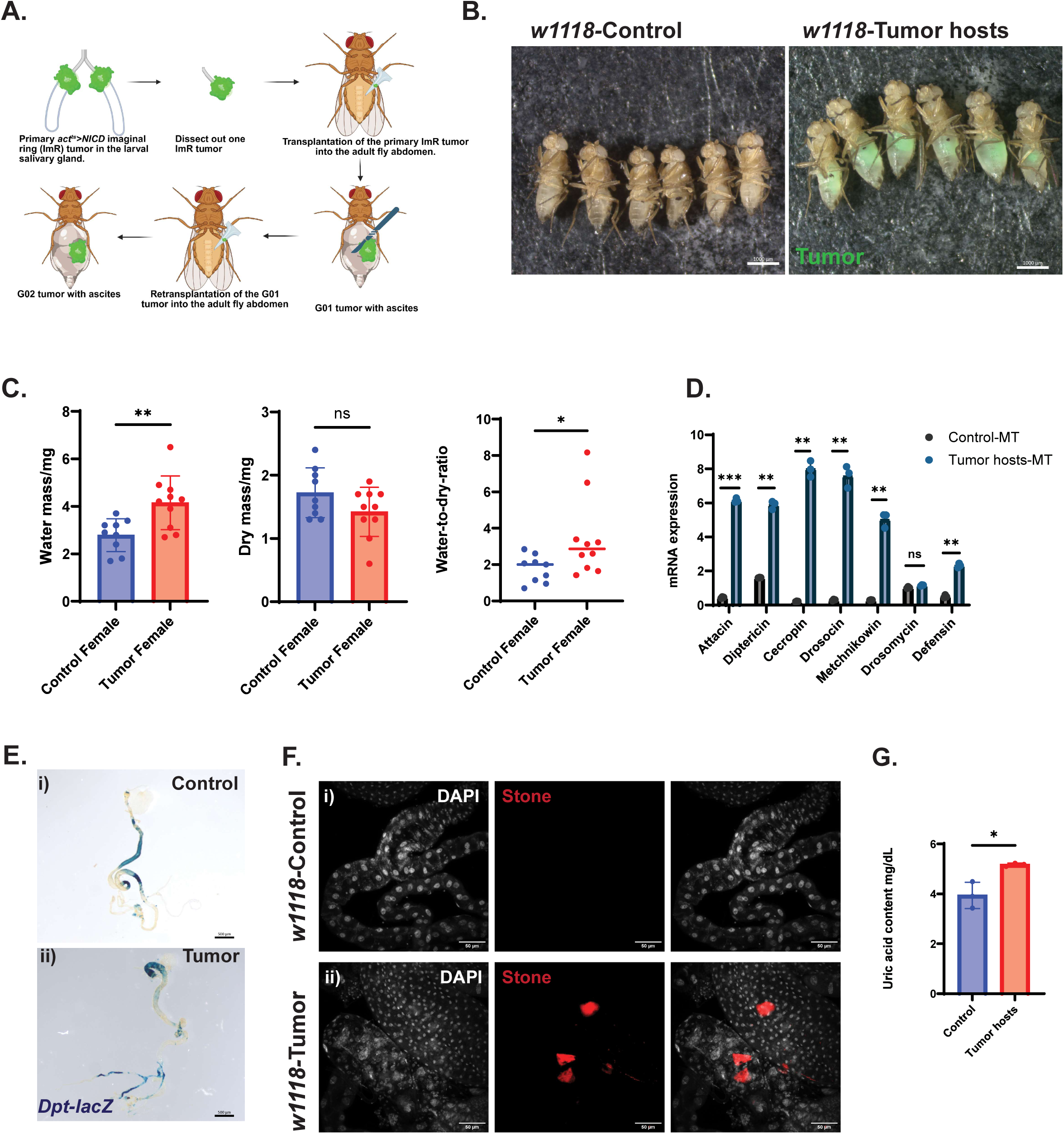
High-passage *act^ts^>NICD* tumor transplantation induces Malpighian tubules-specific innate immune activation, kidney stone formation, and ascites development. **A.** The procedure for allografting *NICD*-overexpressed salivary gland transition zone tumor -*act-Gal4, tub-Gal80>UAS-NICD, CD8GFP* (*act^ts^>NICD)-* into the adult fly abdomen. *act^ts^>NICD* tumors were dissected from larvae in Schneider’s (SD) medium for transplantation. Flies injected with primary *act^ts^ >NICD* tumors (Generation 0-G0) were regarded as Generation 01-G01 hosts. After resting at room temperature for 1 day, the G01 flies were transferred to 29°C to promote tumor tissue growth. To pass the tumor tissue, the G01 tumor was harvested from the host body 9-10 days after initial tumor transplantation and was cut into small pieces, each as large as the primary tumor. These small G1 tumor pieces were then transplanted into new adult flies, which then became the G02 tumor hosts. After passing through several generations (continuous re-transplantation of the same tumor) in the adult fly abdomen, the tumor is called a high passage tumor (HPT). **B.** Abdominal bloating of the 9-day-old wild-type *w^1118^* HPT hosts reared at a 29°C incubator with a 12h/12h light/dark cycle (right). 9-day-old control flies were mock-injected with SD medium (left) and maintained in the same condition as tumor-bearing hosts. **C.** Water mass, dry mass, and water-to-dry mass ratio of the 9-day-old *w^1118^* control and HPT hosts. Each dot represents each biological replicate. Each replicate consisted of 4 whole adult flies. The significance was tested by performing the unpaired t-tests. **D.** The mRNA expression of the antimicrobial peptides of the Malpighian tubules (MTs) tissue collected from 9-day-old *w^1118^* SD-injected control and HPT hosts. Each dot represents biological replicates. Each replicate consisted of 42 pairs of MTs. The significance was tested by performing multiple paired t-tests. **E. β**-Galactosidase (β-Gal) staining of the gut and MTs of the 9-day-old *Diptericin-lacZ* reporter-SD injected i) control and ii) HPT hosts. Scale bar: 500 µm. **F.** MTs’ lower segment region of the 9-day-old i) control and ii) HPT hosts showing kidney stones (red) in tumor-bearing hosts. The nucleus was stained with DAPI (white). Scale bar = 50µm. **G.** The uric acid content of the 9-day-old *w^1118^* whole adult flies: SD injected control and HPT hosts. Each dot represents biological replicates. Each replicate consisted of 4 adult flies. All the flies were female reared at a 29°C incubator with a 12h/12h light/dark cycle. Data are presented as means ± SEM. *p<0.05, **p<0.01, ***0<0.001. ****p<0.0001, ns=non-significant.

HPT TZ tumors were previously shown to compromise gut barrier integrity, leading to bacterial translocation and innate immune activation in host nephrocytes (16). To determine whether renal tubules are similarly affected, we examined immune activation in the Malpighian tubules (MTs), of tumor-bearing hosts. Quantitative RT–PCR analysis revealed robust upregulation of multiple antimicrobial peptide (AMP) genes in MTs from tumor-bearing flies. In *Drosophila*, the IMD and Toll pathways constitute the major humoral immune defenses: the IMD pathway primarily targets Gram-negative bacteria, whereas the Toll pathway defends against Gram-positive bacteria and fungi (18).

Seven representative AMPs were examined, including *Attacin, Cecropin,* and *Metchnikowin* (co-regulated by Toll and IMD); *Diptericin and Drosocin* (**I**MD-specific); and *Drosomycin and Defensin* (Toll-specific). IMD-responsive AMPs showed the most prominent induction in tumor hosts, including *Attacin, Cecropin, Metchnikowin, Diptericin,* and *Drosocin*. Although the Toll pathway AMP *Defensin* was also elevated, the antifungal AMP *Drosomycin* remained unchanged (Fig. 1D). This expression profile indicates predominant activation of the IMD/NF-κB pathway in MTs. MT-specific IMD/NF-κB activation was independently confirmed using *Diptericin-lacZ* reporter flies, which displayed strong β-Galactosidase (β-Gal) staining in the MTs of tumor-bearing hosts but not in renal tubules of SD-injected controls (Fig. 1E). Together, these findings demonstrate that HPT tumors trigger robust renal tubule innate immune activation.

In parallel with IMD/NF-κB activation, kidney stone-like structures became evident in the lower tubules of tumor-bearing hosts (Fig. 1F). IMD activation in *Drosophila* MTs enhances purine metabolism (19, 20), thereby increasing uric acid production. In humans, elevated uric acid levels in blood (hyperuricemia) or urine (hyperuricosuria) are established risk factors for gout and kidney stone formation (21). Consistent with these observations, total uric acid levels were significantly elevated in tumor-bearing hosts compared with SD-injected controls (Fig. 1G). These findings link tumor-induced renal tubule immune activation to purine metabolic dysregulation and nephrolithiasis.

Together, these observations establish that HPT tumors induce systemic fluid accumulation accompanied by IMD/NF-κB dependent renal tubule immune and metabolic pathology.

### Innate immune suppression rescues tumor-associated ascites

To determine whether IMD activation contributes to ascites development in tumor-bearing flies, we tested whether reducing gut bacteria could alleviate fluid accumulation in tumor-bearing hosts. Because IMD signaling is stimulated by gut bacterial overgrowth and translocation into the hemolymph (16), we employed two bacterial depletion strategies: (i) feeding hosts 5 µM tert-butyl hydroperoxide (tBH) and (ii) feeding hosts a cocktail of antibiotics. Early-life exposure to low-dose tert-butyl hydroperoxide (tBH) has been shown to eliminate the gut bacterium *Acetobacter* and extends lifespan (19). Consistent with this, we found that feeding tumor-bearing hosts 5 µM tBH reduced expression of Relish (the IMD/NF-κB transcriptional activator) in the fat body (Sl. Fig. 1A, B). Antibiotic treatment, in contrast, broadly eliminates gut bacteria (16, 20). Both tBH and antibiotic feeding markedly suppressed abdominal fluid accumulation in HPT-bearing hosts (Fig. 2A). A similar rescue was observed in 21-day-old flies injected with *retn>NICD* primary tumors and maintained on either tBH- or antibiotic-containing diets at 25 °C (Sl. Fig. 1C, D).

**Fig. 2:**
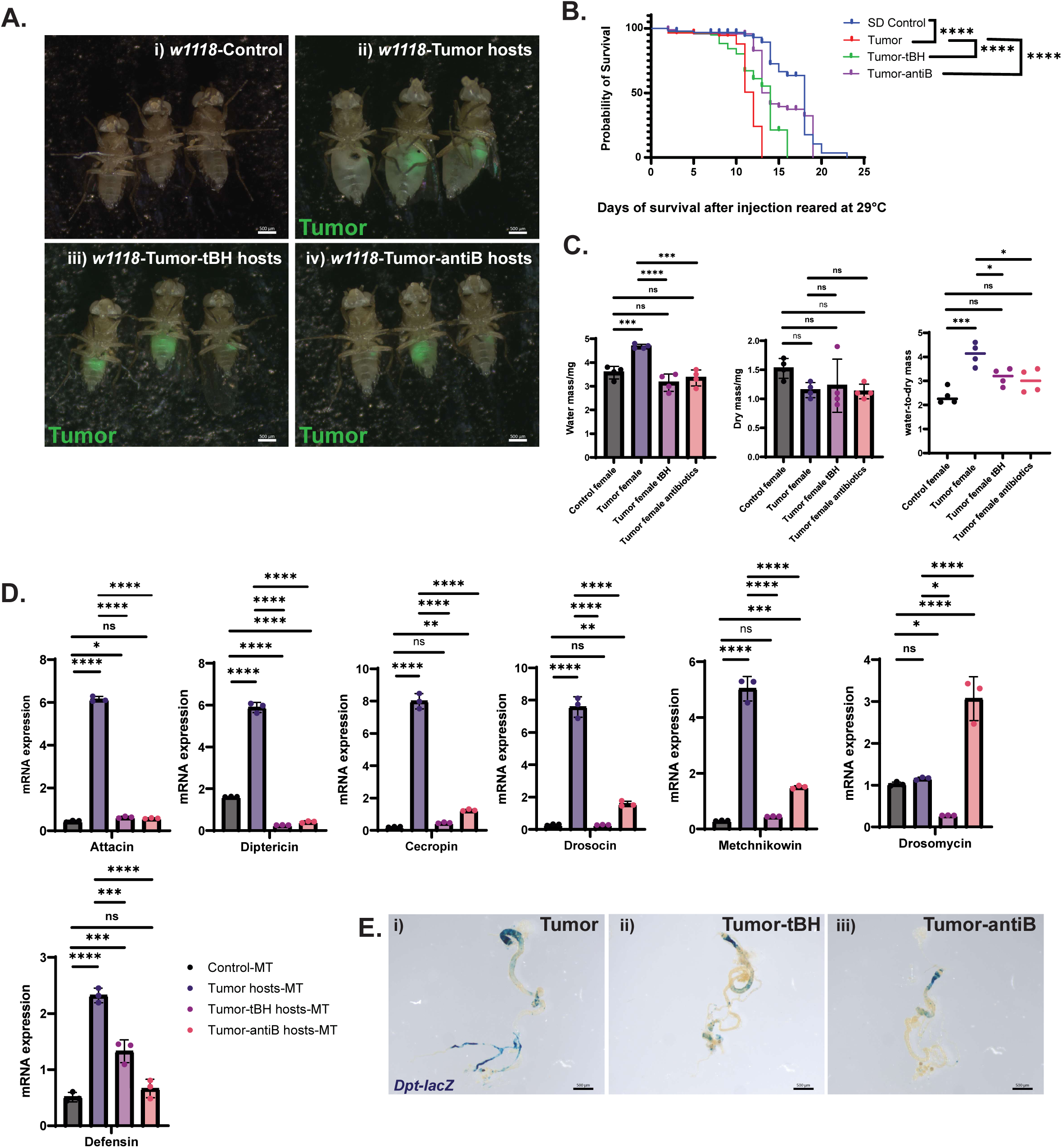
Removal of gut bacteria, particularly *Acetobacter,* attenuates ascites development and Malpighian tubules-specific IMD/NF-kB activation in tumor-bearing hosts. **A.** The image of the 9-day-old *w^1118^* i) Schneider’s (SD) medium injected control, ii) high-passage tumor (HPT) allografted hosts reared in regular cornmeal food, iii) HPT hosts reared in 5µm tert butyl hydroperoxide (tBH) containing cornmeal, and iv) HPT hosts reared in antibiotics containing cornmeal. All the flies rested at room temperature for 1 day after injection and then transferred and maintained at 29°C incubator with a 12h/12h light and dark cycle till Day 9. **B.** Survival graph of the control (n=27) and HPT hosts reared in a regular cornmeal diet (n=44), and tBH (n=43) and antibiotics (n=36) containing cornmeal diet. The significance of the survival data was measured with the Kaplan-Meier survival analysis. **C.** Water mass, dry mass, and water-to-dry mass ratio of the 9-day-old *w^1118^* control and HPT host flies reared in regular, tBH, and antibiotics-containing diet. Each dot represents each biological replicate. Each replicate consisted of 4 whole adult flies. **D.** The mRNA expression of the antimicrobial peptides of the Malpighian tubules (MTs) tissue collected from 9-day-old *w^1118^* SD-injected control and HPT hosts reared in regular, tbH, and antibiotics-containing cornmeal diet. Each dot represents biological replicates. Each replicate consisted of 42 pairs of MTs. **E.** β-Galactosidase (β-Gal) staining of the gut and MTs of the 9-day-old *Diptericin-lacZ* reporter-i) HPT hosts in regular, ii) HPT hosts in tBH, and iii) HPT hosts in antibiotics containing diet. All the flies were female reared at a 29°C incubator with a 12h/12h light/dark cycle. Data are presented as means ± SEM. *p<0.05, **p<0.01, ***0<0.001. ****p<0.0001, ns=non-significant. The significance was tested by performing the one-way ANOVA with Tukey post hoc test.

Consistent with improved physiological status, both tBH and abtibiotics treatments significantly improved survival of tumor-bearing hosts (Fig. 2B). Total body water content was significantly reduced in tBH- or antibiotic-fed tumor hosts, whereas dry weight remained comparable to that of tumor-bearing flies maintained on regular food. Consequently, the water-to-dry weight ratio was decreased in treated animals (Fig. 2C). These results indicate that bacterial depletion ameliorates tumor-induced systemic fluid imbalance.

Because bacterial depletion alleviated ascites, we next examined its impact on renal tubule immune activation. MTs isolated from tumor hosts reared on tBH or antibiotic diets displayed significant reductions in mRNA levels of multiple AMPs, including *Attacin, Diptericin, Cecropin, Drosocin, Metchnikowin,* and *Defensin*, whereas Drosomycin was unchanged (Fig. 2D). This pattern is consistent with selective attenuation of IMD pathway activity. Notably, Drosomycin transcripts were elevated in antibiotic-fed hosts, potentially reflecting fungal expansion following broad bacterial depletion.

Attenuation of MT-specific IMD/NF-κB activation in treated tumor hosts was independently confirmed using the Diptericin-lacZ reporter (Fig. 2E). β-Gal staining showed minimal differences in the gut among tumor-bearing flies fed regular, tBH-, or antibiotic-containing diets. In contrast, reporter activity was markedly reduced in the MTs of treated tumor hosts. Together, these findings indicate that tBH responsive gut bacteria are required for tumor-associated renal tubule IMD activation and ascites development.

### Bacterial depletion suppresses tumor-induced uric acid accumulation and metabolic dysfunction in the MTs

Because bacterial depletion attenuated ascites and renal tubule IMD activation, we next asked whether these interventions also mitigate tumor-associated nephrolithiasis and purine metabolic dysregulation. Tumor-bearing flies fed either tBH- or antibiotic displayed a marked reduction in kidney stone burden, reflected by decreases in both stone size and stone frequency (Fig. 3A). Stone incidence declined from 73% (16/22) in tumor hosts on a regular diet to 35% (8/23) and 25% (6/24) in the tBH- and antibiotic-treated groups, respectively.

**Fig. 3:**
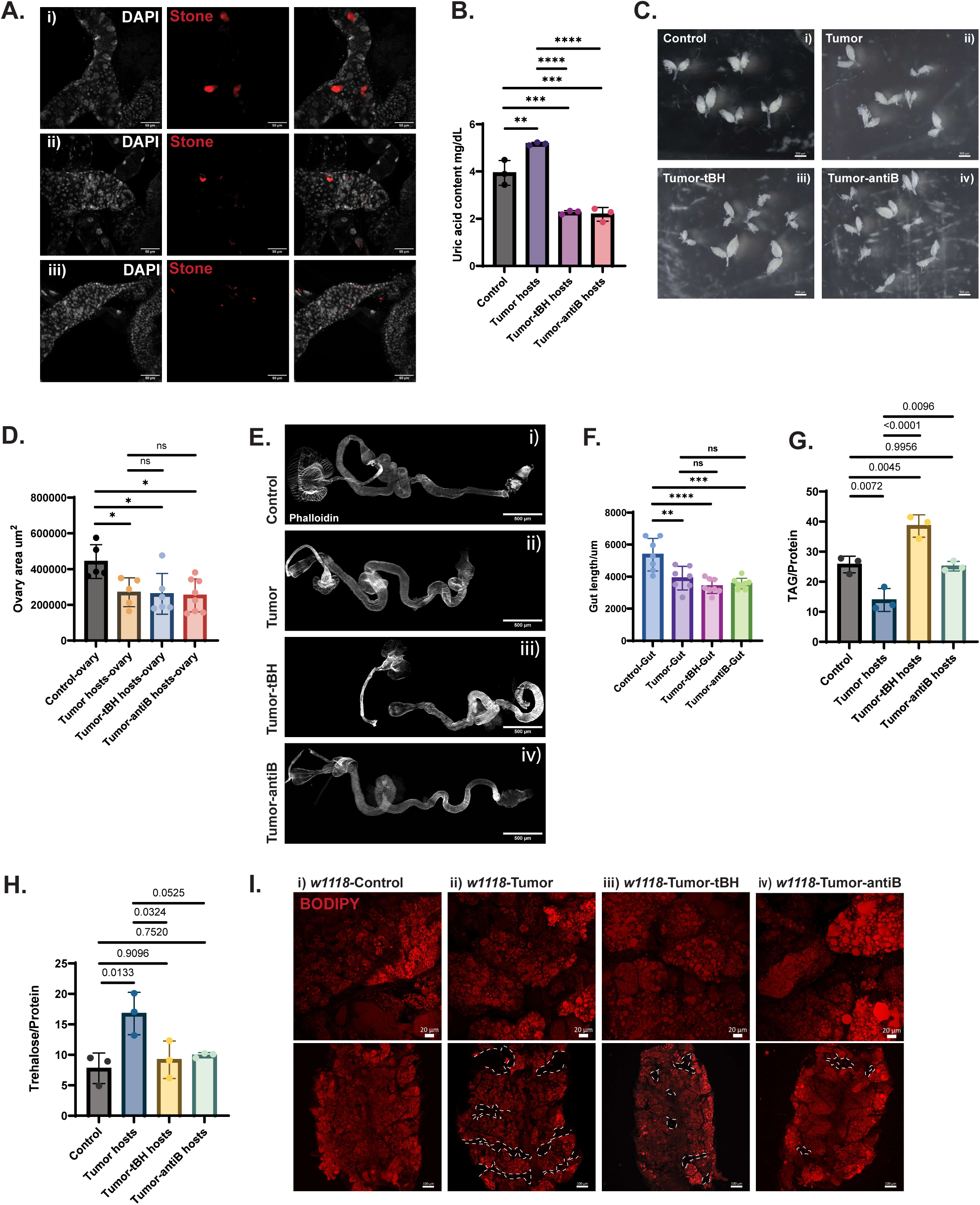
Attenuation of the gut bacterial expansion reduces kidney stone formation and rescues the metabolic imbalance of the tumor-bearing flies. **A.** Malpighian tubules’ (MTs) lower segment region of the 9-day-old *w^1118^* high-passage tumor (HPT) hosts in i) regular, ii) tBH, and iii) antibiotic containing cornmeal diet, showing kidney stone (red). The dissolution of the kidney stones was observed in the tBH and antibiotics-fed HPT hosts’ MTs. The nucleus was stained with DAPI (white). Scale bar = 50µm. **B.** The quantification of the uric acid content in the whole adult flies. Four groups are 9-day-old *w^1118^* Schneider’s (SD) medium injected control, HPT hosts reared in regular, tBH, and antibiotic-containing cornmeal diet. Each dot represents biological replicates. Each replicate consisted of 4 adult flies. **C.** The image of the ovaries of the 9-day-old *w^1118^ -* i) SD injected control, HPT allografted tumor hosts in ii) regular, iii) tBH, and iv) antibiotics containing cornmeal diet. Scale bar = 500µm. **D.** The surface area of the ovary was measured using ImageJ. **E.** The gut of the 9-day-old *w^1118^* -i) SD injected control, HPT allografted tumor hosts in ii) regular, iii) tBH, and iv) antibiotics-containing food was stained with Phalloidin. Scale bar = 500µm. **F.** The gut length was measured using ImageJ. **G.** Triglyceride (TAG) content in the whole body of the 9-day-old *w^1118^ -*SD injected control, HPT allografted tumor hosts in regular, tBH, and antibiotics-containing food. The TAG content was normalized with the total protein content. **H.** Trehalose content in the hemolymph of the 9-day-old *w^1118^ -*SD injected control, HPT allografted tumor hosts in regular, tBH, and antibiotics-containing food. The trehalose content was normalized with the total protein content. **I.** Lipid droplets in the fat body of the 9-day-old *w^1118^ –* i) SD injected control, HPT allografted tumor hosts in ii) regular, iii) tBH, and iv) antibiotics-containing food were stained with BODIPY™. The loss of fat in the HPT hosts was drawn with dotted lines. Scale bar = 20µm and 200 µm. All the flies were female reared at a 29°C incubator with a 12h/12h light/dark cycle. Data are presented as means ± SEM. *p<0.05, **p<0.01, ***0<0.001. ****p<0.0001, ns=non-significant. The significance of the statistical analysis was measured with the one-way ANOVA with Tukey post hoc test.

HPT hosts also displayed elevated uric acid accumulation. Consistent with suppression of IMD-driven purine metabolism, dietary administration of either tBH or antibiotics markedly reduced whole-body uric acid levels, paralleling the decrease in stone formation (Fig. 3B). These findings further support a functional link between microbiota-dependent immune activation and tumor-associated purine metabolic remodeling.

HPT hosts exhibit multiple cachexia-like phenotypes, including reduced gut length, ovarian atrophy, decreased triglyceride storage, and elevated circulating trehalose (16). To determine whether bacterial depletion broadly improves tumor-associated metabolic dysfunction, HPT-bearing flies were fed either tBH or antibiotics. Neither treatment rescued ovarian surface area (Fig. 3 C,D), or gut length (Fig. 3E, F), indicating that these wasting phenotypes are largely microbiota independent.

In contrast, both interventions markedly increased triglyceride storage (Fig. 3G), while reducing circulating trehalose levels (Fig. 3H) in host flies. These results indicate that microbiota-dependent inflammation contributes selectively to systemic metabolic imbalance in tumor hosts.

Lipid staining of fat bodies further supported this conclusion. Tumor-bearing hosts exhibited enlarged lipid droplets relative to controls but showed an overall reduction in fat body tissue (dotted outlines). Tumor-bearing flies fed tBH- or antibiotic-containing diets likewise displayed enlarged lipid droplets, but the reduction in fat body tissue was less pronounced than in hosts maintained on a regular diet (Fig. 3H). Together, these findings indicate that bacterial depletion partially restores metabolic homeostasis in tumor-bearing flies.

### Tumors induce species-specific gut dysbiosis that amplifies IMD signaling

We next asked whether tumor-associated immune activation arises from defined changes in gut microbial composition. To determine whether tumor-bearing hosts develop gut dysbiosis and whether bacterial depletion strategies reverse this microbial overgrowth, we cultured homogenized gut tissues from 9-day-old flies on MRS plates. Culture for 2 days at 30°C revealed a marked increase in colony-forming units (CFUs) in tumor-bearing hosts compared with controls (Fig. 4A), consistent with microbial overgrowth and dysbiosis(19). We next examined the impact of bacterial depletion strategies on this phenotype. Rearing tumor-bearing flies on either tBH or an antibiotic-containing diet significantly reduced gut CFUs. tBH treatment lowered CFU counts consistent with selective depletion of *Acetobacter* species, whereas antibiotic feeding eliminated detectable bacterial growth (Fig. 4A). These results confirm that tumor hosts develop microbiota-dependent dysbiosis that is reversible by bacterial depletion.

**Fig. 4:**
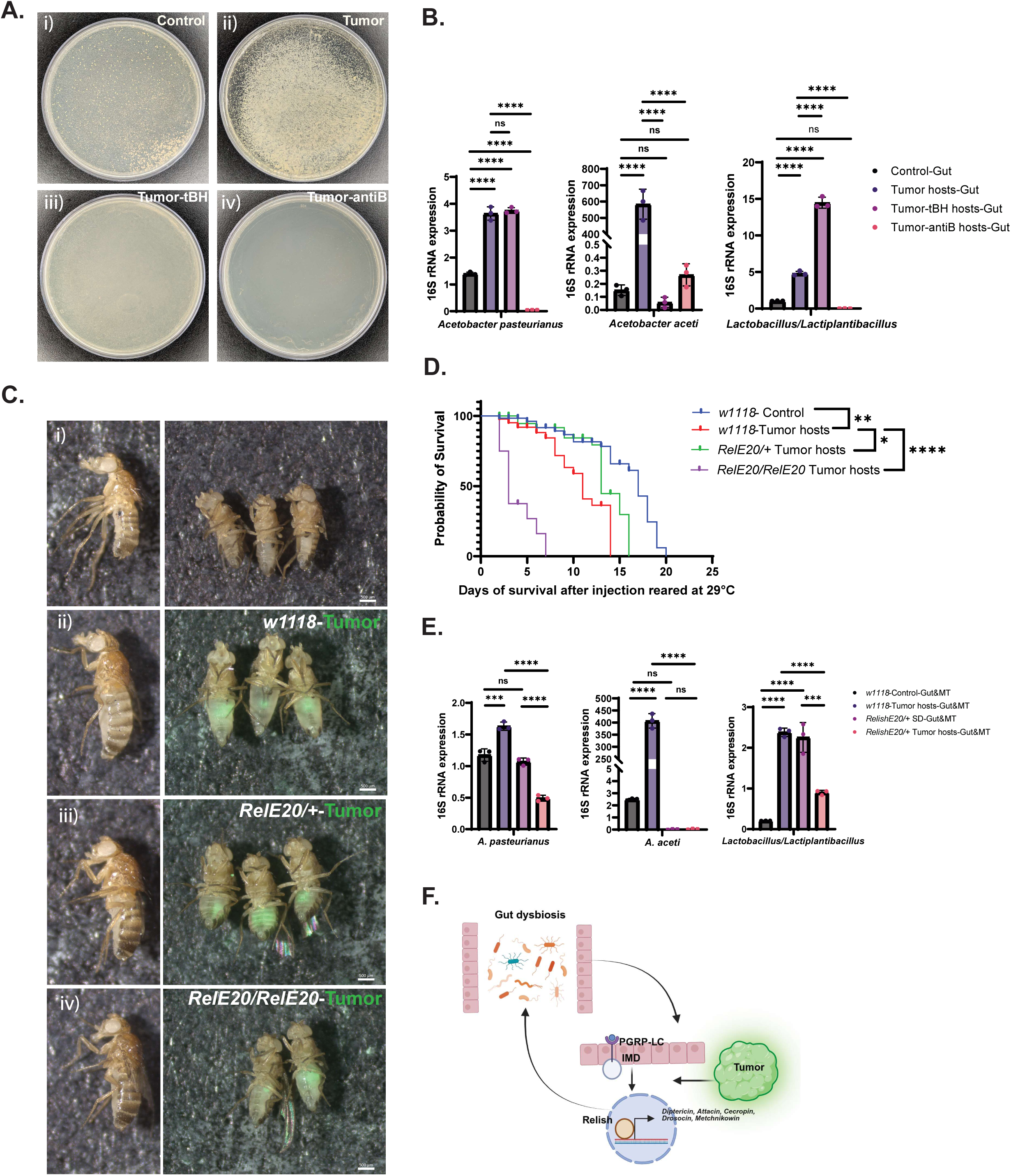
Tumor-induced expansion of the Acetobacter aceti causes IMD/NF-kB activation. **A.** De Man–Rogosa–Sharpe agar plating of the homogenized gut tissues collected from 9-day-old *w^1118^* i) Schneider’s (SD) medium injected control, High-passage tumor (HPT) allografted female hosts in ii) regular, iii) tBH, and iv) antibiotics-containing food. **B.** The quantitative real-time PCR of the 16S ribosomal RNA of the bacteria residing in the 9-day-old *w^1118-^*SD medium injected control, HPT allografted hosts in regular, tBH, and antibiotics-containing food. The statistical significance of the graph was measured using the paired t-test **C.** The image of the 9-day-old -i) *w^1118^* SD injected control, ii) *w^1118^* HPT allografted hosts, iii) heterozygous *Relish* mutant *Rel^E20^/+* HPT allografted hosts, and iv) homozygous *Relish* mutant *Rel^E20^/Rel^E20^* HPT allografted hosts. Scale bar = 500µm. **D.** Survival graph of the *w^1118^* SD injected control (n=22), *w^1118^* HPT allografted hosts (n=17), heterozygous *Relish* mutant *Rel^E20^/+* HPT allografted hosts (n=19), and homozygous *Relish* mutant *Rel^E20^/Rel^E20^*HPT allografted hosts (n=22). The significance of the survival was measured with the Kaplan-Meier survival analysis. **E.** The quantitative real-time PCR of the 16S ribosomal RNA of the gut bacteria residing in the 9-day-old *w^1118^ -*SD injected control, *w^1118^* HPT allografted hosts, heterozygous *Relish* mutant *Rel^E20^/+ -* SD injected control, and heterozygous *Relish* mutant *Rel^E20^/+* HPT allografted hosts. The statistical significance of the graph was measured using the paired t-test. **F.** The illustration shows tumor-induced IMD activation and microbial expansion form a positive feedback loop. All the flies were female reared at a 29°C incubator with a 12h/12h light/dark cycle. Data are presented as means ± SEM. *p<0.05, **p<0.01, ***0<0.001. ****p<0.0001, ns=non-significant. The significance of the statistical analysis was measured with the one-way ANOVA with Tukey post hoc test.

Previous 16S sequencing analyses revealed that both control and tumor-bearing hosts harbor two predominant bacterial genera, *Acetobacter* and *Lactiplantibacillus* (16). Among these, *Acetobacter* is dominant and comprises two species in the fly gut, *Acetobacter pasteurianus* and *Acetobacter aceti*.

To determine whether tumor-associated dysbiosis is species resolved, we quantified individual bacterial species by qRT–PCR targeting 16S rRNA. Both *Acetobacter* species and *Lactobacillus* were elevated in tumor host guts. Strikingly, *A. aceti* exhibited an approximately 3000-fold increase relative to SD-injected controls (Fig. 4B), whereas *A. pasteurianus* showed only a modest ∼2.5-fold elevation, and *Lactobacillus/Lactiplantibacillus* increased ∼5-fold (Fig. 4B). These data identify *A. aceti* as the dominant species selectively expanded in tumor hosts.

Although prior work reported that tBH eliminates the entire *Acetobacter* genus (20), quantitative RT–PCR analysis of gut tissue from tumor hosts fed 5 µM tBH revealed selective effects: *A. aceti* was effectively eliminated, whereas *A. pasteurianus* levels remained largely unchanged. Thus, tBH enables functional dissection of species-specific contributions to tumor-associated pathology. Because tBH treatment both extends tumor host lifespan and reduces abdominal fluid accumulation, the selective loss of *A. aceti* suggested that this species may disproportionately contribute to IMD activation, and ascites, a possibility we tested directly below.

We next investigated how tumors promote dysbiosis. Previous work showed that constitutive activation of the IMD pathway in enterocytes or expression of the Toll ligand Spätzle in the fat body increases the abundance of *Gluconobacter* (21). These findings raised the possibility that tumor-induced IMD activation and microbial expansion form a positive feedback loop.

To test this model, HPTs were allografted into homozygous and heterozygous *Relish* mutant flies-*Rel^E20^*. Relish is the transcription factor of the IMD pathway (22), and its nuclear translocation is required for AMP production. Neither homozygous *Rel^E20^/Rel^E20^*nor heterozygous *Rel^E20^/+* tumor hosts developed abdominal fluid accumulation following tumor transplantation (Fig. 4C). However, homozygous *Rel^E20^/Rel^E20^* tumor-bearing hosts exhibited reduced lifespan compared with wild-type *w^1118^* tumor hosts (Fig. 4D), whereas heterozygous *Rel^E20^/+* tumor-bearing hosts showed improved survival (Fig. 4D).

To assess gut microbial abundance in this sensitized background, we quantified *A. pasteurianus*, *A. aceti*, and *Lactobacillus* in *Rel^E20^/+* control and tumor hosts. Abundance of both *Acetobacter* and *Lactobacillus* species was significantly reduced in *Rel^E20^/+* tumor hosts relative to wild-type *w^1118^* tumor hosts (Fig. 4E). Partial reduction of IMD signaling significantly suppressed tumor-associated bacterial expansion, with the most pronounced effect observed for *A. aceti* (Fig. 4E). Together, these findings support a positive feedback loop in which *Relish*/IMD/NF-κB signaling promotes tumor-associated species-specific dysbiosis that further amplifies systemic immune signaling (Fig. 4F).

### *A. aceti* is necessary and sufficient for tumor-associated IMD activation and ascites

Mono-association of defined bacterial species in *Drosophila* has been widely used to interrogate microbial contributions to host physiology (23, 24). Because tBH treatment selectively eliminates *A. aceti* while sparing *A. pasteurianus* and concurrently alleviates ascites in tumor-bearing hosts, we hypothesized that expansion of *A. aceti* is sufficient to drive the ascitic phenotype.

To test this possibility, axenic (germ-free) flies were generated and mono-associated with either *A. pasteurianus* or *A. aceti* (Fig. 5A). Individual *Acetobacter* species were isolated from tumor-bearing fly guts by culturing homogenates on MRS agar. Colonies were distinguished based on morphology: *A. pasteurianus* forming smooth colonies and *A. aceti* exhibiting a spiral appearance (Fig. 5B), and species identity was confirmed by PCR amplification and Sanger sequencing of the bacterial 16S rDNA region. HPTs were then allografted into axenic and mono-associated flies. Because axenic and gnotobiotic flies display reduced physiological robustness relative to conventional hosts, animals were maintained at 29 °C for 6 days and then shifted to 25 °C for 5 days prior to analysis. Under these conditions, ascites developed selectively in *A. aceti* mono-associated tumor hosts, whereas axenic flies and those mono-associated with *A. pasteurianus* showed no abdominal bloating despite persistent tumor growth (Fig. 5C). Similar results were obtained using the *retn>NICD* primary tumor model maintained at 25 °C for 21 days (Sl. Fig. 1E). confirming the species-specific effect.

**Fig. 5:**
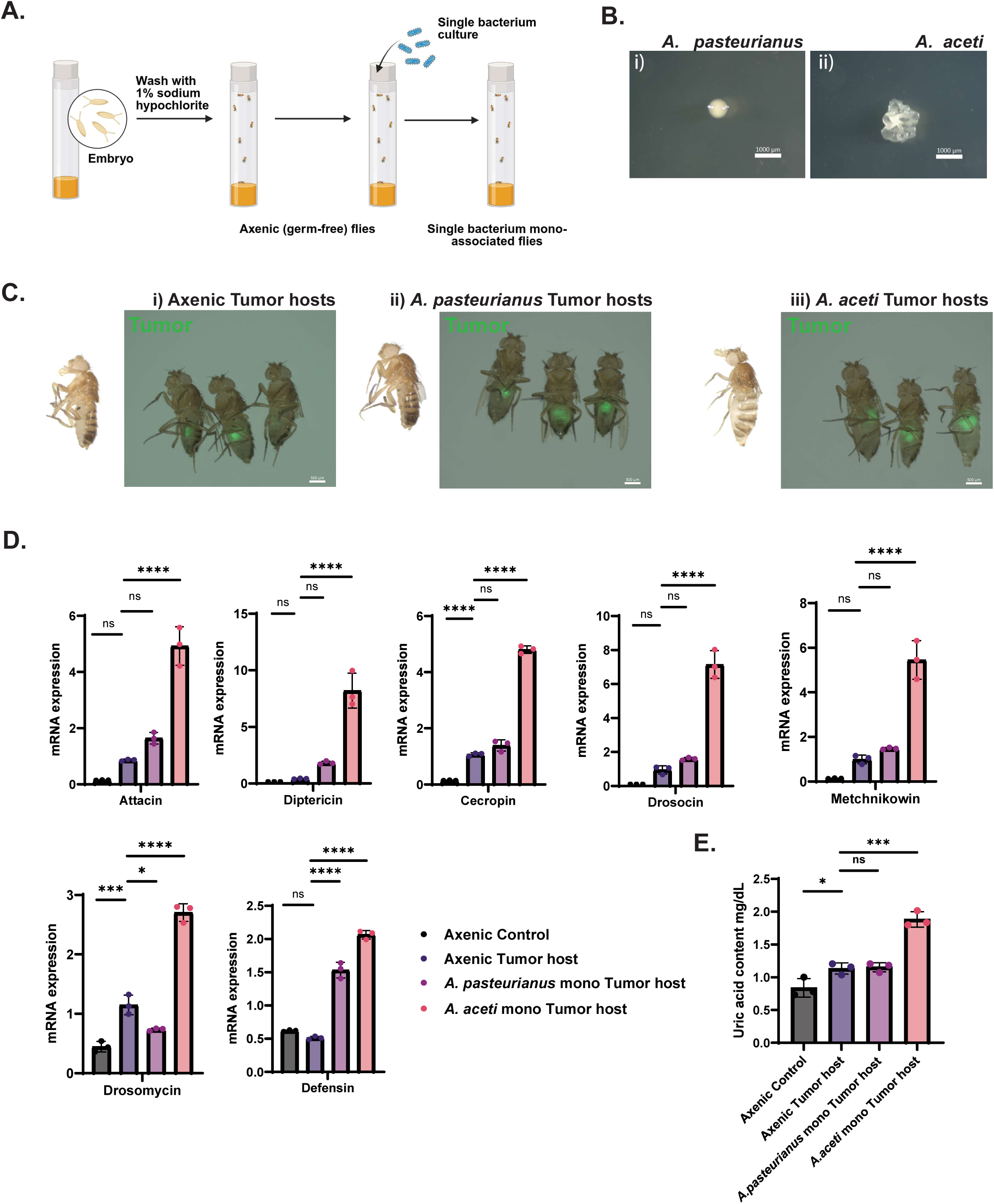
Mono-association of *Acetobacter aceti* confirms their sole contribution in tumor-induced ascites development. **A.** The illustration of developing axenic (germ-free) flies and mono-association of the axenic flies with *Acetobacter* species. First, the embryos collected were washed with 1% sodium hypochlorite to dechorionate the embryos. Then the embryos were kept in sterile autoclave fly food and maintained at a 25°C incubator. The adult flies that emerged were germ-free. For bacterial mono-association, 150 μl of bacterial culture (OD_600_ =1) was applied to the autoclaved fly food containing germ-free embryos and maintained at a 25°C incubator. The eclosed adult flies now contained a single bacterial species. **B.** The single colony of the bacteria i) *Acetobacter pasteurianus* and ii) *Acetobacter aceti*. Scale bar = 1000µm. **C.** The image of the 11-day-old *w^1118^* – i) axenic high-passage tumor (HPT) hosts, ii) *A. pasteurianus* mono-associated HPT hosts, and iii) *A. aceti* mono-associated HPT hosts, reared at 29°C for 6 days and 25°C for 5 days. **D.** The global mRNA expression of the antimicrobial peptides of the 9-day-old *w^1118^* -SD injected axenic control and axenic HPT hosts, *A. pasteurianus,* and *A. aceti* mono-associated HPT hosts, reared at a 29°C incubator with a 12h/12h light/dark cycle. Tumor tissues were removed prior to the hosts’ tissues collected for the RNA extraction. **E.** The uric acid content of the 9-day-old *w^1118^* whole adult flies: SD injected axenic control, HPT allografted axenic hosts, *A. pasteurianus,* and *A. aceti* mono-associated HPT hosts. Each dot represents biological replicates. Each replicate consisted of 4 adult flies. All the flies were female. Data are presented as means ± SEM. *p<0.05, **p<0.01, ***0<0.001. ****p<0.0001, ns = no significance. The significance of the statistical analysis was measured with the one-way ANOVA with Tukey post hoc test.

To determine whether differential immune activation underlies these phenotypes, the expression of global antimicrobial peptides (AMPs) was quantified. Tumor-bearing axenic flies mostly showed no significant elevation in AMP expression relative to SD-injected axenic controls. Exceptions were *Cecropin* and *Drosomycin*, which exhibited upregulation in the axenic tumor hosts. Compared to axenic tumor hosts, tumor-bearing flies associated with *A. pasteurianus* exhibited only minor ∼3-fold increases in *Defensin* expression and marked downregulation in *Drosomycin* expression. Other AMPs showed no significant change. However, mono-association with bacteria *A. aceti* amplified this response. *A. aceti* association produced a robust ∼5–10-fold increase of IMD-responsive AMPs (Fig. 5D), indicating that *A. aceti* is a dominant contributor to IMD/NF-κB activation in tumor-bearing flies.

Because IMD signaling has been linked to purine metabolism (24), we next tested whether *A. aceti*–driven immune activation correlates with uric acid production. Consistent with a causal role, uric acid levels were significantly elevated only in tumor hosts harboring *A. aceti*, but not in axenic hosts or those associated with *A. pasteurianus* showed no significant change (Fig. 5E).

Together, these findings demonstrate that *A. aceti* is both necessary and sufficient to drive tumor-associated IMD activation, purine metabolic dysregulation, and ascites formation.

### MT-intrinsic IMD activation is sufficient and necessary to drive ascites

To determine whether IMD pathway activation within Malpighian tubules (MTs) is sufficient to drive renal tubule pathology, we selectively activated IMD signaling in MT principal cells. Overexpression of PGRP-LC, the receptor that recognizes Gram-negative peptidoglycan (22), or Relish, under the *CG31272-Gal4* driver, which expressed in principal cells of the stem cell zone and main segment of the upper tubules (Sl. Fig. 2A). Consistently, pronounced ascites-like fluid accumulation was induced even in the absence of tumors by day 4 at 29 °C (Fig. 6A). Thus, MT-restricted IMD activation is sufficient to trigger systemic fluid retention. These findings are consistent with recent work demonstrating that constitutive IMD/NF-κB activation in MT principal cells is sufficient to trigger abdominal fluid retention (25).

**Fig. 6:**
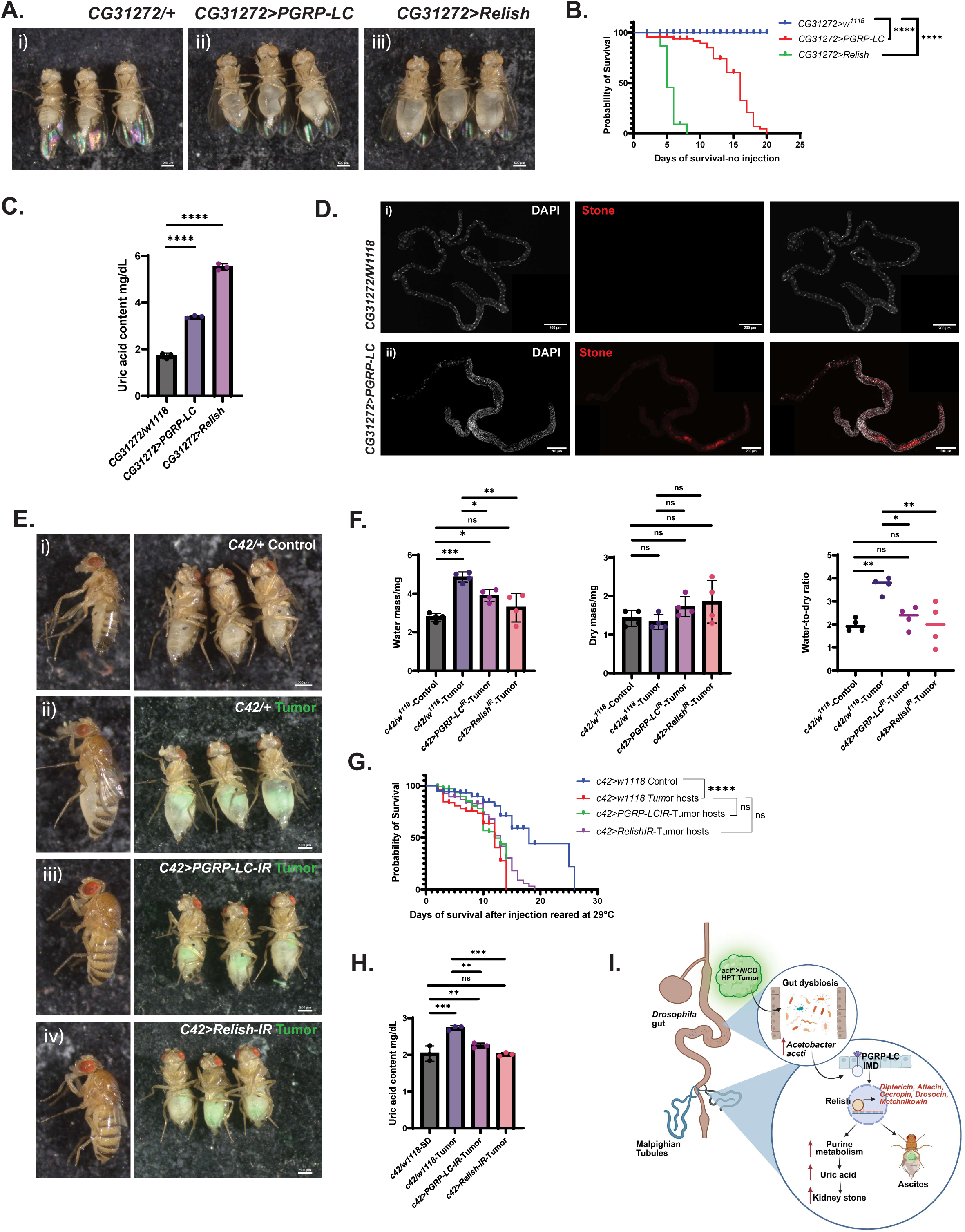
Suppression of the Malpighian tubules-specific IMD/NF-kB activation rescues ascites and reduces uric acid content of the tumor-bearing flies. **A.** The image of the 4-day-old i) *CG31272/+*, ii) *CG31272>PGRP-LC*, and iii) *CG31272>Relish,* shows abdominal fluid accumulation with the constitutive expression of the *PGRP-LC,* and *Relish* expressed under the Malpighian tubules (MTs) Gal4. B. Survival graph of the female *CG31272/+* (n=30)*, CG31272>PGRP-LC* (n=49), and *CG31272>Relish* (n=21) flies. The significance of the survival was measured with the Kaplan-Meier survival analysis. C. The uric acid content of the 4-day-old whole adult flies-*CG31272/+, CG31272>PGRP-LC*, and *CG31272>Relish*. Each dot represents biological replicates. Each replicate consisted of 4 adult flies. D. The image of the Malpighian tubules (MTs) of the 7-day-old i) *CG31272/+* control and ii) *CG31272>PGRP-LC* flies, showing kidney stones (red) in the MTs of the *CG31272>PGRP-LC* flies. The nucleus was stained with DAPI (white). Scale bar = 200µm. E. The image of the 9-day-old i) *C42/+* SD injected control, ii) *C42/+* HPT allografted hosts, iii) *C42>PGRP-LC^RNAi^* HPT hosts, and iv) *C42>Relish ^RNAi^* HPT hosts. Scale bar = 500µm. F. Water mass, dry mass, and water-to-dry mass ratio of the 9-day-old *C42/+* SD injected control, *C42/+* HPT allografted hosts, *C42>PGRP-LC ^RNAi^* HPT hosts, and *C42>Relish ^RNAi^* HPT hosts. Each dot represents each biological replicate. Each replicate consisted of 4 whole adult flies. G. Survival graph of the *C42/+* SD injected control (n=38), *C42/+* HPT allografted hosts (n=69), *C42>PGRP-LC ^RNAi^* HPT hosts (n=48), and *C42>Relish ^RNAi^* HPT hosts (n=77). The significance of the survival data was measured with the Kaplan-Meier survival analysis. H. The uric acid content of the 9-day-old whole adult flies-*C42/+* SD injected control, *C42/+* HPT allografted hosts, *C42>PGRP-LC ^RNAi^* HPT hosts, and *C42>Relish ^RNAi^* HPT hosts. Each dot represents biological replicates. Each replicate consisted of 4 adult flies. I. The schematic diagram shows how tumors induce gut dysbiosis, where the expansion of the *Acetobacter aceti* activates the IMD/NF-kB signaling in the tumor-bearing hosts’ Malpighian tubules. The activated IMD/NF-kB signaling enhances purine metabolism, upregulates uric acid production, and kidney stone formation. Simultaneously, activated IMD/NF-kB signaling causes ascites development. All the flies were female reared at a 29°C incubator with a 12h/12h light/dark cycle. Data are presented as means ± SEM. *p<0.05, **p<0.01, ***0<0.001. ****p<0.0001, ns=non-significant. The significance of the statistical analysis was measured with the one-way ANOVA with Tukey post hoc test.

Consistent with severe physiological stress, MT-specific overexpression of either PGRP-LC or Relish significantly shortened lifespan (Fig. 6B). Because elevated IMD/NF-κB activity in MTs has been linked to purine metabolic remodeling (26), we next examined whether MT-intrinsic IMD activation alters uric acid homeostasis. Whole-body uric acid levels were markedly increased following MT-specific overexpression of PGRP-LC or Relish (Fig. 6C). Moreover, 7-day-old flies overexpressing PGRP-LC in MTs developed widespread kidney stone–like deposits throughout the tubules (Fig. 6D). Owing to reduced survival, stone analysis in Relish-overexpressing flies at day 7 was not feasible; however, stone formation was already evident by day 4 under the same conditions (Sl. Fig. 2B). These results indicate that MT-intrinsic IMD activation is sufficient to drive purine dysregulation and nephrolithiasis.

We next asked whether attenuating IMD/NF-κB signaling in MTs could mitigate tumor-associated pathology. Suppression of IMD signaling in MT principal cells using the *C42-Gal4* driver, which is expressed principal cells throughout the tubules except the ureter (Sl. Fig. 2A) or *CG31272-Gal4* substantially rescued abdominal distention in tumor-bearing hosts (Fig. 6E, Sl. Fig. 2E) and substantially reduced the water-to-dry weight ratio (Fig. 6F). Similar improvement was observed in *retn>NICD* tumor hosts (Sl. Fig. 2C, D), with independent MT-specific RNAi lines targeting *Relish* or *PGRP-LC* in hosts. These findings indicate that MT-intrinsic IMD activity is required for tumor-associated fluid accumulation.

While MT-specific suppression of IMD signaling through knockdown of *PGRP-LC* or *Relish* in the renal tubules did not significantly improve survival of tumor-bearing flies (Fig. 6G), it substantially reduced systemic uric acid accumulation (Fig. 6H). Together, these results position renal IMD/NF-κB signaling as the proximal effector of tumor-induced purine dysregulation, nephrolithiasis, and ascites.

## Discussion

The relationship between host health and gut microbiota is complex and multifaceted. Chronic metabolic conditions, including obesity, type 2 diabetes, fatty liver disease, and cardiometabolic disorders, have been associated with alterations in gut microbial composition (26). Modulation of the microbiota has also been explored therapeutically in neurological disorders such as autism spectrum disorder, Alzheimer’s disease (27–29), Parkinson’s disease (30–33), attention deficiency hypersensitivity disorders, and depression (34–36). Despite these advances, the contribution of specific microbial species to tumor-associated renal dysfunction and fluid imbalance remains poorly defined. In the present study, we identify a species-resolved microbiome mechanism that promotes renal pathology and ascites in tumor-bearing *Drosophila*, underscoring the importance of defining microbiota–host interactions that may contribute to nephrogenic ascites in cancer.

Here, we show that expansion of *A. aceti* in tumor-bearing flies drives systemic IMD/NF-κB activation. In response to bacterial challenge, flies mount an innate immune response characterized by increased antimicrobial peptide (AMP) production. *Drosophila* immune defenses comprise four major signaling arms: IMD and Toll pathways, which primarily regulate AMP production, and the JNK and JAK/STAT pathways. The JNK pathway responds broadly to cellular stressors—including reactive oxygen species, UV exposure, DNA damage, infection, and tissue injury (37). whereas JAK/STAT signaling enhances stress resistance and promotes expression of targets such as Turandot A.

Previous work in the *esg^ts^>yki*^3SA^ gut tumor model demonstrated elevated JNK/Jra signaling in MT principal cells and showed that expression of dominant-negative Basket (the *Drosophila* JNK homolog) partially rescues abdominal bloating (13). In another *esg^ts^>yki^3SA^* gut tumor model study, an antidiuretic hormone, ion transport peptide isoform F (ITP_F_), was shown to act on stellate cells via TkR99D to inhibit fluid secretion (14).

Independent studies in a *Drosophila* ovarian cancer model reported that tumor-derived Upd2 activates JAK/STAT signaling in MT stellate cells and secondarily stimulates ERK/MAPK activity in renal stem cells (RSCs), driving hyperproliferation and fluid accumulation (15). Our findings define a mechanistically distinct pathway in which species-specific gut dysbiosis engages MT-intrinsic IMD/NF-κB signaling to promote nephrogenic ascites. Consistent with a role for innate immune signaling in MT remodeling, forced activation of PGRP-LC or Relish in MTs in our study induced increased cell proliferation within the stem cell zone of the lower tubules even in the absence of tumors (Fig. 6D, Sl. Fig. 2B). These findings suggest that IMD/NF-κB activity may contribute to RSC hyperproliferation observed in tumor-associated renal dysfunction.

The divergence among tumor models likely reflects differences in how tumor-derived signals engage host physiology. In the current model, *NICD-*driven tumorigenesis is confined exclusively to the transplanted tumor tissue. In contrast, in gut and ovarian tumor models, the host organs themselves are genetically manipulated, e.g., *yorkie* overexpression in the gut or *Ras^V12^* and *aPKC^ΔN^* overexpression in the ovary, directly altering host tissues. We propose that these distinct upstream triggers converge on shared downstream renal effectors controlling fluid balance, including ion transporters, water channels, or purine metabolic pathways. Future work will be required to define the precise transport mechanisms affected in Malpighian tubules during tumor-associated fluid imbalance

Using the allografted high-passage *act^ts^>NICD* tumor model, we demonstrate that MT-specific IMD activation is a key driver of ascites-like fluid accumulation. We further show that systemic IMD activation in tumor-bearing flies originates from gut dysbiosis, specifically from overexpansion of *A. aceti*. Tumor growth is associated with chronic inflammation that perturbs microbial homeostasis, and partial reduction of *Relish* activity normalizes bacterial overabundance in tumor hosts. These data support a feed-forward loop in which tumor-induced inflammation promotes species-specific dysbiosis that further amplifies systemic immune activation. The resulting expansion of *A. aceti* further amplifies IMD signaling, establishing a self-reinforcing inflammatory loop that promotes purine metabolic remodeling and uric acid accumulation. Consequently, tumor hosts develop kidney stone–like deposits in the MTs alongside ascites. Both phenotypes are suppressed either by selective elimination of *A. aceti* or by attenuation of IMD signaling within MTs.

Although prior studies have linked IMD activation to purine metabolism (26, 38), the molecular mechanisms connecting innate immune signaling to metabolic rewiring remain incompletely defined. Our data place MT-intrinsic IMD/NF-κB signaling as the proximal effector linking microbiota-dependent inflammation to nephrogenic ascites. While previous work proposed an association between nephrolithiasis and abdominal fluid retention (13), establishing a unified mechanistic framework will require further investigation. In summary, this study identifies species-specific expansion of the Gram-negative commensal *A. aceti* as a key upstream driver of tumor-associated renal dysfunction and ascites in *Drosophila*. These findings reveal a tumor–microbiome–immune–renal axis that couples microbial imbalance to systemic fluid homeostasis and suggest that targeting defined microbial species may represent a strategy to mitigate cancer-associated renal and fluid complications.

## Materials and Methods

### *Drosophila* stock maintenance

The details of the fly stocks and genotypes used for each experiment are listed in *SI Appendix, Table S1*. All the flies collected for each experiment were reared in the same environment, and the experiments were performed at the same time to avoid batch effects.

### Fly Feeding

The regular diet contains fly food recipe by Blooming Drosophila stock center – BDSC Cornmeal food. Tert-Butyl hydroperoxide food contained 5ml of the 100 µM of the tBH solution, which was mixed with 100ml of BDSC Cornmeal food. Antibiotic cocktail food contained 15mg Carbenicillin, 15mg Metronidazole, and 7.5mg Tetracycline mixed with 100 ml of BDSC Cornmeal food.

### Tumor transplantation

All the tumor-allografted female flies and Schneider’s medium (SD) injected control flies were virgin and collected right after eclosion. The flies were kept at room temperature for 24 hours prior to injection. All female-bearing hosts carried high-passage tumors that were allografted according to the protocol described by Cong et al (16). Briefly, the primary *act^ts^>NICD* tumors were dissected from larvae in Schneider’s (SD) medium for transplantation. Flies injected with primary *act^ts^ >NICD* tumors (G0) were regarded as G01 hosts. After resting at room temperature for 1 day, the G01 flies were transferred to a 29°C incubator with a 12h/12h light and dark cycle to promote tumor tissue growth. To pass the tumor tissue, the G01 tumor was harvested from the host body 9-10 days after initial tumor transplantation and was cut into small pieces, each as large as the primary tumor. These small G01 tumor pieces were then transplanted into new adult flies, which then became the G02 tumor hosts. After the tumor passes through several generations (continuous re-transplantation of the same tumor) in the adult fly abdomen, the tumor is now called a high passage tumor (HPT). The *retn>NICD* tumor closely resembles the *act^ts^>NICD* tumor, with two key differences. First, *retn-Gal4* restricts *NICD* overexpression specifically to the posterior transition zone of the salivary gland, leading to neoplastic tumor growth, while the hyperplastic anterior expansion observed in the *act^ts^>NICD* model is absent. Second, *Gal4* activity in this model is not regulated by *tubulin-Gal80*, allowing *NICD* expression to occur independently of temperature (17). Thus, flies injected with *retn>*NICD tumor were reared at a 25°C incubator with a 12h/12h light and dark cycle.

### Survival quantification

After tumor transplantation or mock injection with Schneider’s (SD) medium, the flies were kept at room temperature for 24 hours. Then, the next day, the flies were kept at 29 °C, and each day the number of flies that died was noted. The food vials were changed every 3 days, so the flies do not die by sticking to the old food. For the tert-Butyl hydroperoxide or antibiotics-treated flies, they were fed the modulated food from the day of injection and continuously throughout their lifespan. After the data were collected, graphs were plotted using the software GraphPad Prism. The significance of the statistical analysis was measured with the log-rank (Mantel-Cox) test.

### Reagents, equipment, and other

The details of the reagents and equipment utilized are listed in *SI Appendix, Table S3*.

### RNA extraction, cDNA synthesis and quantitative real time PCR

Tissues were collected in 100 μl TRIzol® Reagent and stored at -80 °C until RNA extraction. Tissues involving whole flies include 4-5 whole flies. For the gut bacterial analysis, 21 gut samples were collected from each group. For the analysis involving the Malpighian tubules tissues 42 pairs of tubules were collected from each group.

RNA extraction is performed using the Directzol® RNA Miniprep kit supplied by Zymo Research, following the manufacturer’s protocol. cDNA syntheses were performed using the Superscript® III First Strand supplied by Invitrogen, following the manufacturer’s protocol.

Quantitative real-time polymerase chain reaction was performed as follows. The reaction mixture containing 1 μl cDNA, 1 μl of 10 μM concentration of forward and reverse primers, and 5 μl iTaq Universal SYBR Green Supermix (Bio-Rad, Cat#172-5124) was loaded onto a 96-well plate. The CFX96 Touch Real-Time PCR System (Bio-Rad) was used for qRT-PCR analysis. Primer sequences used for qPCR are provided in (*SI Appendix, Table S2*). All the primers, except 16S ribosomal RNA primers, were designed using Primer-BLAST (https://www.ncbi.nlm.nih.gov/tools/primer-blast) and manufactured by Eurofins Genomics (eurofinsgenomics.com). The sequence for the 16S ribosomal RNA primers for *A. pasteurianus* and *A. aceti* were adapted from the study of Torija et al (39) and the primer sequence for *Lactobacillus/Lactiplantibacillus* was adapted from the study of Shi et al (40). Glyceraldehyde-3-phosphate dehydrogenase (GAPDH) and ribosomal protein 49 (rp49) mRNA were used for normalization.

### Fly water mass and dry mass measurement

The mass of the flies was measured at the age of Day 9 of the tumor transplantation. Each sample (each dot in the graph) contains 4 flies. At least four independent biological replicates were cultivated for each group. The wet mass of each sample was measured in the analytical balance. After measuring wet mass, the flies were kept at -80°C to freeze their tissue. Then the frozen samples were desiccated in Vacufuge® plus with vacuum aqueous (V-AQ) mode for two hours. The desiccated samples were measured again to quantify the dry mass of the flies. Water content in each sample was measured by subtracting the wet mass from the dry mass.

### Uric acid quantification

Four flies were collected in the 100 µl lysis buffer (0.2% Triton X-100 in 1X PBS) for the uric acid quantification in each sample. The samples were then homogenized and centrifuged at 16000 g for 2-3 mins. 5µl of the supernatant was collected to perform uric acid quantification using the QuantiChrom™ Uric Acid Assay Kit (Cat. No. DIUA-250), following the manufacturer’s protocol. Uric acid was quantified at the wavelength OD_590nm_. Each group consisted of three independent biological replicates.

### TAG, Trehalose quantification

Four to five whole flies were collected in 200µl of the lysis buffer (1x PBS and 0.2% Triton), and the tissue was homogenized and incubated at 70°C for 5 mins. The homogenized sample was then centrifuged at 16000g for 3 mins, and then the supernatant was collected and stored at - 80°C until further downstream application. Triglyceride storage was quantified using the Abcam Triglyceride Assay Kit, utilizing the manufacturer’s protocol, with some modifications. Briefly, for the triglyceride assay, tumors were removed prior to the tissue collection, as triglycerides (TAG) stored in the tumor tissue might have impacted the result for TAG storage of the hosts. Free glycerol was quantified first by incubating the samples with reaction mix for 50 minutes at room temperature. Free glycerol was first measured and then 2µl of cholesterol esterase were added and the sample mixture was incubated at room temperature for 10-15 mins prior quantification. The enzyme will break down the stored TAG into glycerol. Free glycerol and stored triglyceride were quantified at wavelength OD_570nm_.

For trehalose quantification, whole flies were utilized without tumor removal prior to tissue collection, as trehalose circulates in the fly hemolymph. Quantification was performed using the Megazyme Trehalose kit, utilizing the manufacturer’s protocol.

Both the TAG and trehalose measurement was normalized with the protein level, quantified with BCA Protein Assay with BSA Protein Standard kit utilizing the manufacturer’s protocol. Protein content was quantified at wavelength OD_562nm._

### Bacterial single colony culture

De Man–Rogosa–Sharpe (MRS) plates were used to culture the gut bacteria of the control and tumor-bearing host flies. MRS plates were prepared with 25.5g of MRS broth, 0.5 ml Tween® 80, 7.5g agar, and up to 500ml of distilled water. The mixture was heated and autoclaved before making culture plates and kept stored at 4°C, under sterile conditions, prior to use. For the bacteria colony culture, three fly guts were dissected in sterile 1XPBS and transferred to 200 μl sterile Schneider’s (SD) medium. The gut tissues were homogenized in the SD medium and 50 μl of homogenate were plated onto MRS agar plates using glass beads. The plates were incubated at 30 °C for 2 days.

### Axenic fly production and Bacterial mono-association

Axenic flies were generated by following the protocol published by Koyle et al (41). Briefly, embryos were collected and dechorionated for 1 min in 1% sodium hypochlorite, then washed twice in 70% ethanol and subsequently with autoclaved distilled water. The embryos were transferred into autoclaved fly food. The axenic (germ-free) status was confirmed by performing quantitative RT-PCR of the 16S rRNA region on homogenates of the adult flies as they eclosed and by plating the homogenates on LB agar plates.

The mono-association of germ-free (GF) flies with bacteria was conducted following the protocol outlined by Storelli et al (42). Briefly, 150 μl of bacterial culture (OD600 =1) was applied to the autoclaved fly food containing germ-free embryos. The emerging larvae were allowed to develop on the contaminated media to the adult stage, after which the adults were transferred to new autoclaved fly food. The descendants of these adult flies are mono-associated flies.

### Immunofluorescence staining and imaging

The samples used for antibody, DAPI, or BODIPY™ 558/568 C12 staining include gut, Malpighian tubules (MTs), and fat body. First, the samples were dissected in 1X PBS and fixed in 4% fix solution for 30 minutes for the fat body and 2 hours for the gut and MTs. The samples were then washed thrice for 15 mins with PBT (PBS with 0.2% Triton X-100). 1X PBS was used in place of PBT when washing fix from the fat body because the PBT wash might dissolve the stored lipids in the fat body.

For fat body staining with BODIPY™ and DAPI, stains were prepared by mixing BODIPY™ (1:500) and DAPI (1:1000) with 1X PBS. For gut and MTs staining with Phalloidin and DAPI, stains were prepared by mixing Phalloidin (1:200) and DAPI (1:1000) with 1X PBS. Tissues were then allowed to be incubated in the staining solution mixture for 2 hours at room temperature. Then stain solution was removed, and samples were washed thrice in 1X PBS to remove the excess stain.

For the antibody staining, after the fixing and washing steps, the fat body samples were blocked with PBTG for 1 hour to prevent non-specific primary antibody binding. Then, the anti-Relish primary antibody was dissolved in 1XPBS at a 1:300 ratio, and the samples were incubated in the antibody solution overnight at 4 °C. After staining overnight with the primary antibody, the samples were washed thrice with 1XPBS to remove excess antibody. Then, a mixture of secondary antibody and DAPI was prepared by mixing secondary antibody (1:400) and DAPI (1:1000) in the 1X PBS solution. The tissues were then incubated with a mixture of secondary antibody and DAPI at room temperature for 2 hours. After secondary antibody and DAPI staining, the excess stains and DAPI were washed out by washing the samples thrice with 1X PBS at room temperature for 15 minutes.

The samples were stored in mounting solution at 4°C until slide preparation and immunofluorescence imaging. The staining images were taken under LSM800 microscope.

### β-Galactosidase (β-Gal) staining

After 9-10 days of tumor allograft or mock injection with Schneider’s (SD) medium, seven to eight *Drosophila* gut and MTs samples were collected in 1X PBS. Then the tissues were fixed in 0.5% glutaraldehyde for 15 minutes. After fixation, the samples were washed in 1X PBS twice for 5 minutes. Fresh staining solution was prepared by mixing 100 µl potassium ferri-cyanide (50 mM), 100 µl potassium ferro-cyanide (50 mM) in 800 µl PBS with MgCl_2_ (2 mM). The mixture was preheated at 37 °C for at least 30 minutes, then 5 µl of X-gal (Amresco, Cat#0428) of concentration 200 mg/ml was added. After the washing step, the samples were incubated with the staining solution at 37 °C for 30 minutes. Then excess stains were washed with PBT twice for 15 minutes. The slides were prepared by mounting samples in 100% glycerol.

## Acknowledgement

We thank the Bloomington Drosophila Stock Center for providing fly stocks. We are especially grateful to Dr. Chun Nin Wong for guidance in establishing axenic (germ-free) and bacteria mono-associated fly models. We also appreciate Dr. Xianfeng Wang, Yi-Chun Huang, and other members of the Deng lab for their valuable discussions and contributions. This study was supported by the NIH R01GM072562, R01CA227789, and Ladies Leukemia League, Inc. 2024-2025 grants.

---

**Sl. Fig. 1:**
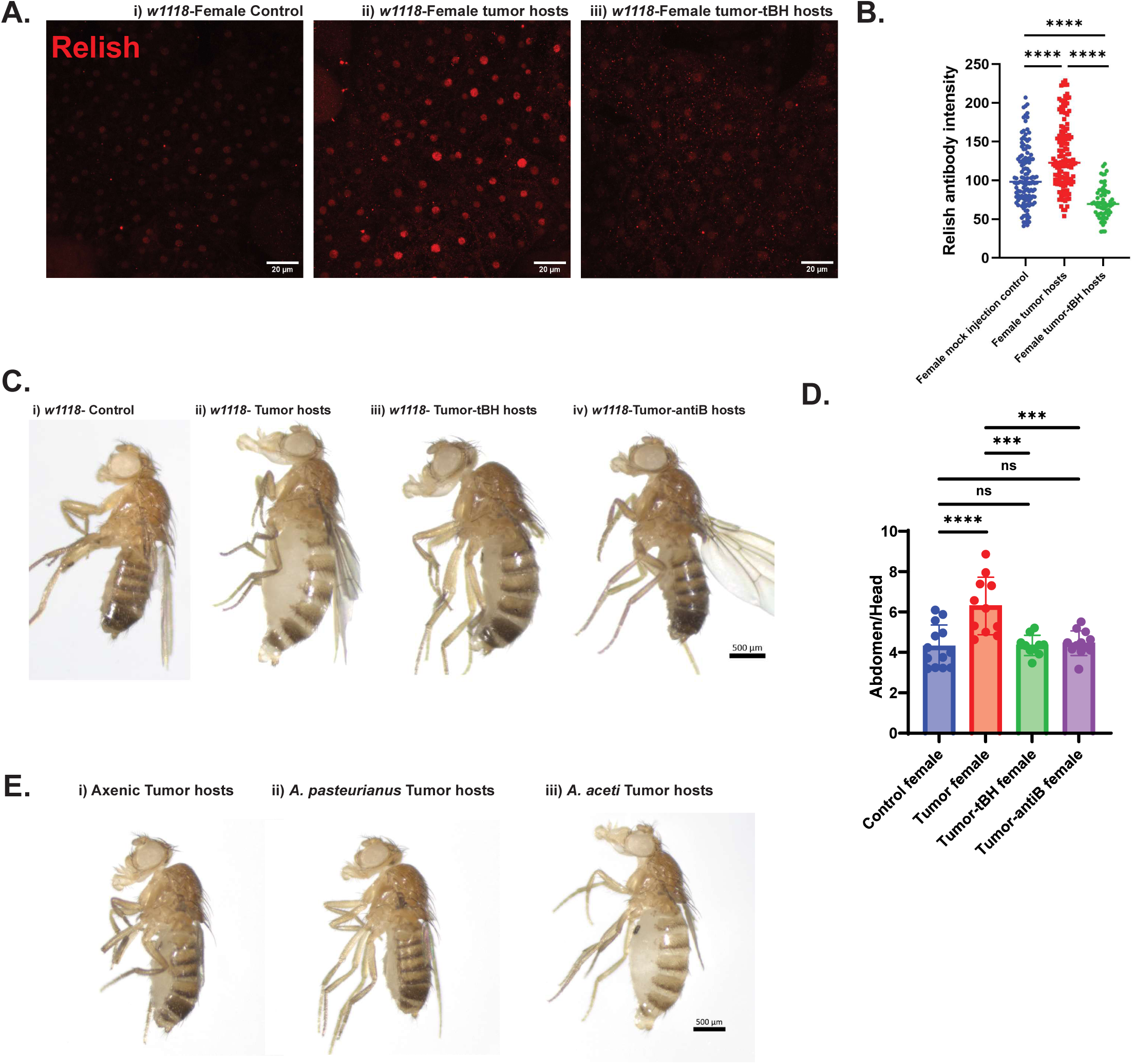
Acetobacter aceti mainly responsible for the tumor induced gut dysbiosis and ascites development. **A.** Fat body of the 21-day-old w1118 i) Schneider’s (SD) medium injected control, ii) primary *retn-Gal4>UAS-NICD,CD8GFP* allografted tumor host-G01 host, and iii) G01 host fed with 5µM tert-butyl hydroperoxide, stained with anti-Relish antibody (red). Scale bar = 20µm. **B.** The signal intensity of the anti-Relish antibody staining in the fat body of the 21-day-old *w1118* SD injected control, primary *retn>NICD* allografted tumor hosts, and tumor hosts fed with 5µM tert-butyl hydroperoxide. The quantification of the signal intensity was performed using ImageJ software. **C.** 28-day-old *w1118* i) SD injected female control flies, ii) primary *retn>NICD* injected tumor host-G01 host, iii) G01 host fed with 5µM tert-butyl hydroperoxide, and iv) G01 host fed with antibiotics. Scale bar = 500µm. **D.** The quantification of the abdomen/head size ratio of the 28-day-old *w1118* SD injected control, primary *retn>NICD* injected tumor host-G01 host, G01 host fed with 5µM tert-butyl hydroperoxide, G01 host fed with antibiotics. **E.** 21-day-old *w1118* i) G01-axenic host, ii) G01-*Acetobacter pasteurianus* mono-associated host, iii) G01-*A. aceti* mono-associated host. All the flies were female reared at a 25°C incubator with a 12h/12h light/dark cycle. Data are presented as means ± SEM. *p<0.05, **p<0.01, ***p<0.001, ****p<0.0001, ns= non-significant. The significance of the statistical analysis was measured with the one-way ANOVA with Tukey post hoc test.

**Sl. Fig. 2:**
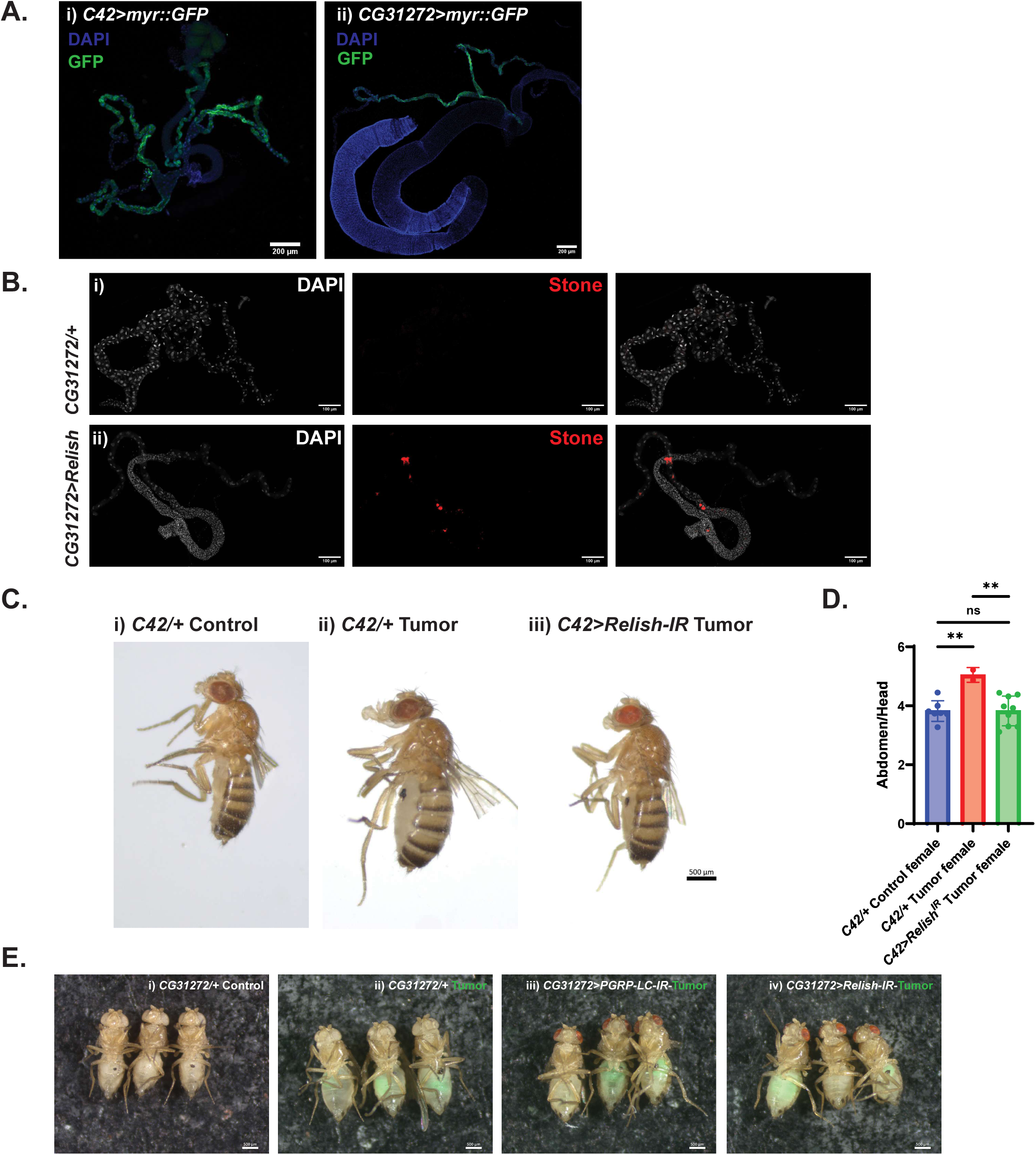
Role of Relish expression in Malpighian tubule principal cells in regulating kidney stone formation and ascites development. **A.** i) *C42>myr::CD8GFP* showing the expression pattern of the *C42-Gal4* in the *Drosophila* Malpighian tubules (MTs). Their expression pattern is all over the MTs’ principal cells, expect the ureter region of the stem cell zones. ii) *CG31272>myr::CD8GFP* showing the expression pattern of the *CG31272-Gal4* in the *Drosophila* MTs. Their expression pattern is from the ureter region till main segment of the MTs. Scale bar = 200µm. **B.** The image of the Malpighian tubules (MTs) of the 4-day-old i) *CG31272/+* control and ii) *CG31272>PGRP-LC* flies, showing kidney stones (red) in the MTs of the *CG31272>Relish* flies. The nucleus was stained with DAPI (white). Scale bar = 100µm. **C.** 28-day-old i) SD injected *C42/+* control, i) primary *retn>NICD* injected *C42/+* tumor host-G01 *C42/+* host, iii) G01 *C42>RelishRNAi* host. Scale bar = 500µm. **D.** The quantification of the abdomen/head size ratio of the 28-day-old female SD injected *C42/+* control, primary *retn>NICD* injected *C42/+* tumor host-G01 *C42/+* host, G01 C42>RelishRNAi host. All the flies were female reared at a 25°C incubator with a 12h/12h light/dark cycle. **E.** The image of the 12-day-old female i) *CG31272/+* SD injected control, high-passage tumor allografted ii) *CG31272/+*, iii) *CG31272>PGRP-LCRNAi*, and iv) *C42>RelishRNAi* hosts, reared at a 29°C incubator with a 12h/12h light/dark cycle. Scale bar = 500µm. Data are presented as means ± SEM. *p<0.05, **p<0.01, ***p<0.001, ****p<0.0001, ns= non-significant. The significance of the statistical analysis was measured with the one-way ANOVA with Tukey post hoc test.

**Sl Appendix, Table S1.**
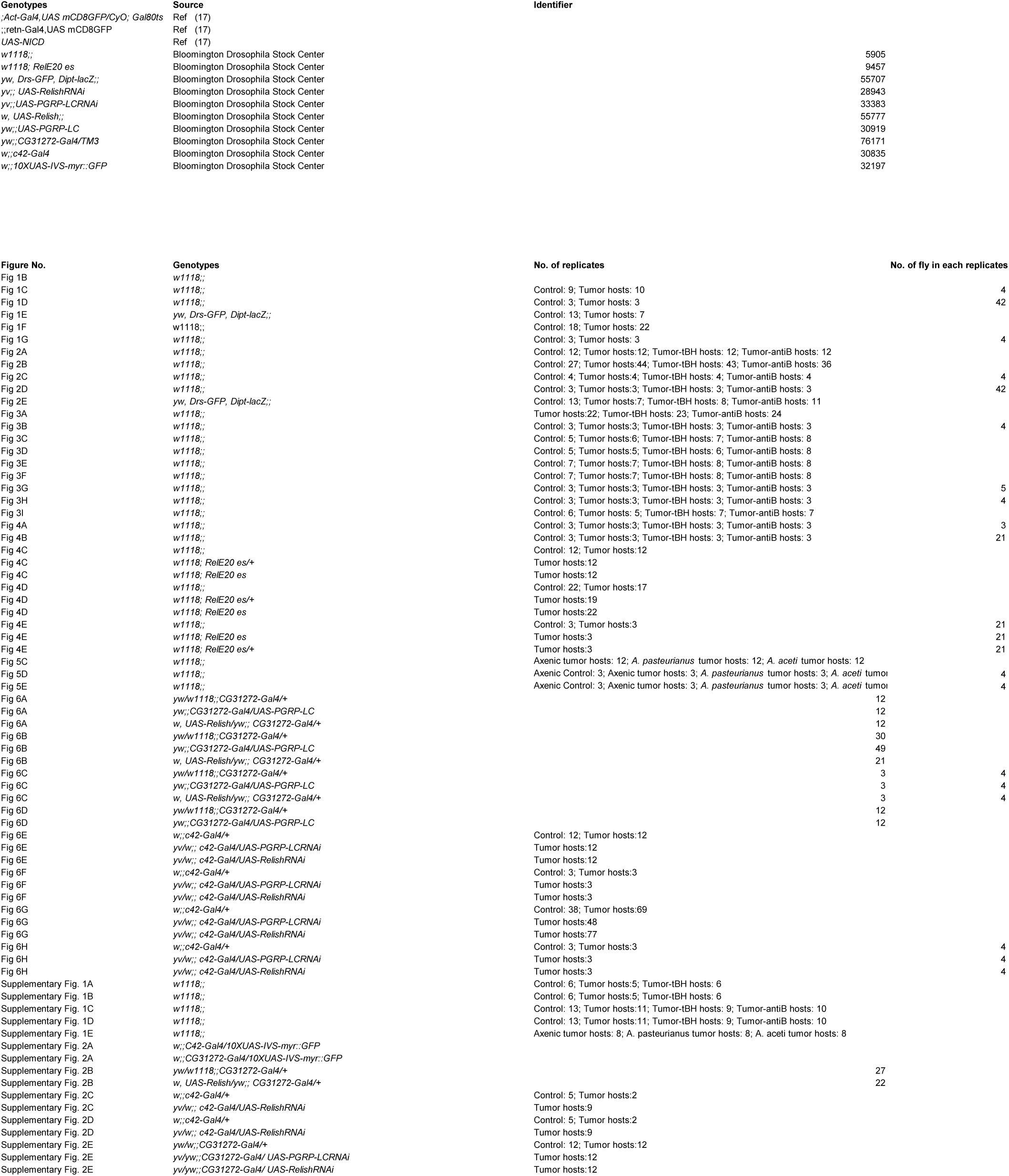
The genotypes, source, identifiers and number of replicates of each fly stock used in the study.

**Sl Appendix, Table S2.**
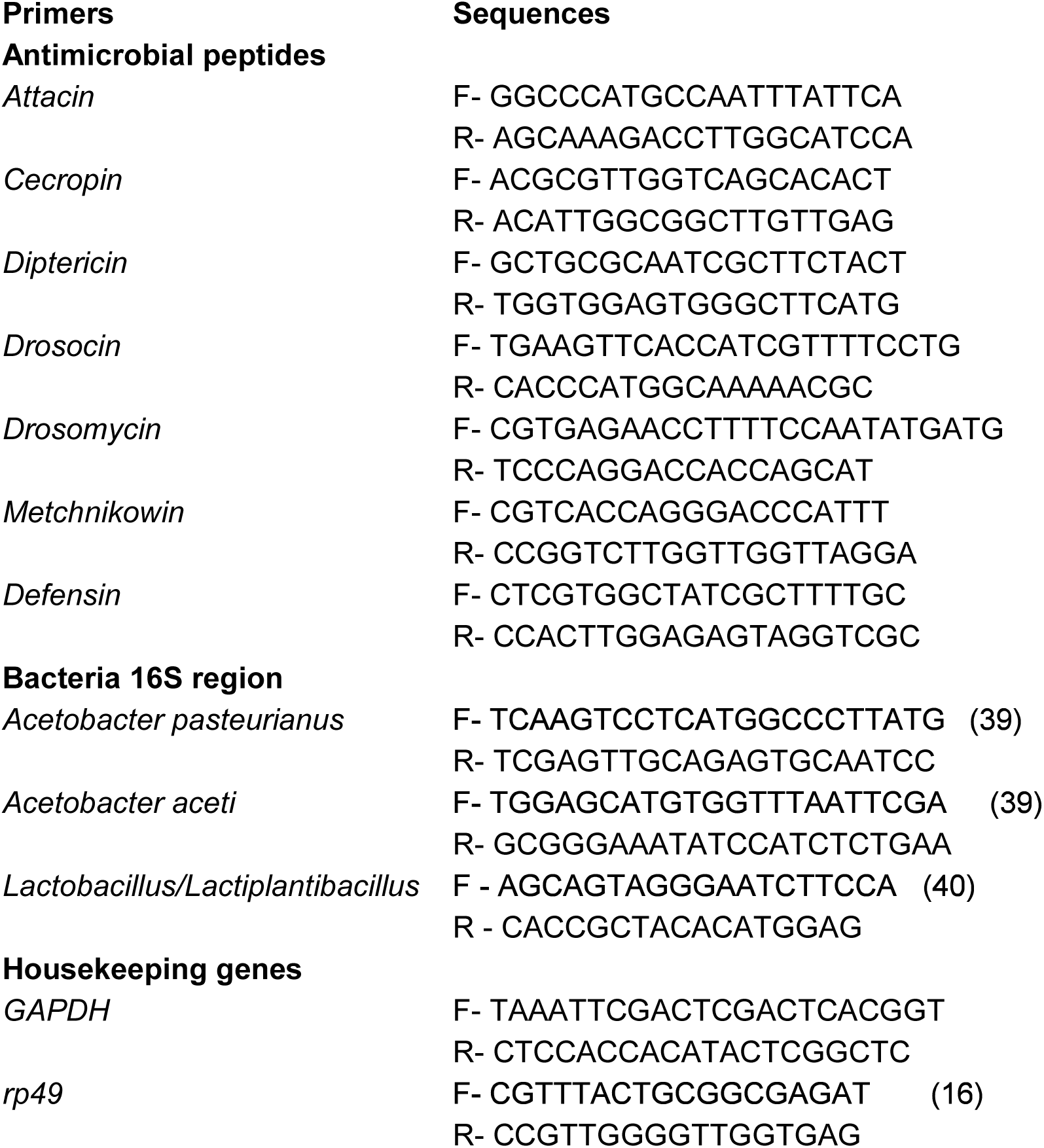
The sequence of the primers used in the study.

**Sl Appendix, Table S3.**
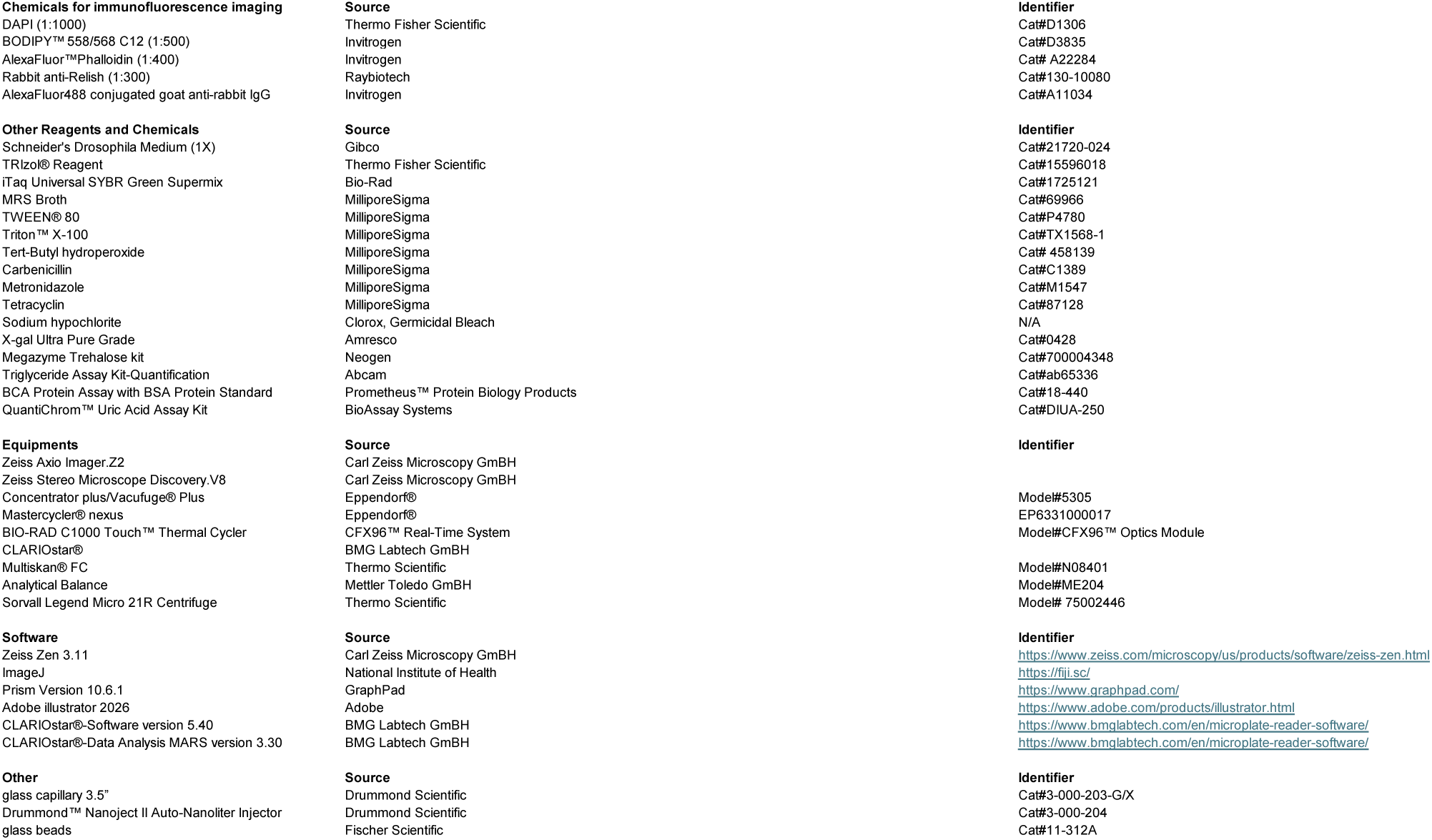
The reagents, chemicals, equipment, softwares used in the study and their sources and identifiers.

## References

1. Goosenberg E, Kudaravalli P, Samant H. Ascites. StatPearls. Treasure Island (FL) 2025.

2. de Moraes FCA, Rego L, Matheus G, Dantas CR, de Tolosa ALM, Burbano RMR, Hirata MH. Does Malignant Ascites Define Prognosis in Gastric Cancer with Peritoneal Spread? A Systematic Review and Meta-analysis. J Gastrointest Cancer. 2025;56(1):206.

3. Jacob R, A S, Abdul Razack N, Prabhuswamimath SC. Malignancy of Malignant Ascites: A Comprehensive Review of Interplay between Biochemical Variables, Tumor Microenvironment and Growth Factors. Asian Pac J Cancer Prev. 2024;25(10):3413–20.

4. Perez-Aizpurua X, Cabello Benavente R, Bueno Serrano G, Alcazar Peral JM, Gomez-Jordana Manas B, Tufet IJJ, et al. Obstructive uropathy: Overview of the pathogenesis, etiology and management of a prevalent cause of acute kidney injury. World J Nephrol. 2024; 13(2):93322.

5. Yousefnezhad A, Yousefi Sharemi SR, Saffarieh E, Nokhostin F. Renal dysfunction in individuals with ovarian cancer; a review on current concepts. J Renal Inj Prev. 2023;12(4): e32247.

6. Muthukumaran A, Wanchoo R, Seshan SV, Gudsoorkar P. Paraneoplastic Glomerular Diseases. Adv Kidney Dis Health. 2024;31 (4):346–57.

7. Amarapurkar P, Bou-Slaiman S, Madrid B, Ladino M. Paraneoplastic glomerular disease: The struggle is real. Journal of Onco-Nephrology. 2019;3(1):31–38.

8. Lee M, Wang Q, Wanchoo R, Eswarappa M, Deshpande P, Sise ME. Chronic Kidney Disease in Cancer Survivors. Adv Chronic Kidney Dis. 2021;28(5):469–76 e1.

9. Sai Spandana G, Viswanathan S, Barathi SD, Selvaraj J. Etiology and Outcomes in Patients With Chronic Kidney Disease and Ascites. Cureus. 2024;16(7):e64113.

10. Ding G, Xiang X, Hu Y, Xiao G, Chen Y, Binari R, et al. Coordination of tumor growth and host wasting by tumor-derived Upd3. Cell Rep. 2021;36(7):109553.

11. Kwon Y, Song W, Droujinine IA, Hu Y, Asara JM, Perrimon N. Systemic organ wasting induced by localized expression of the secreted insulin/IGF antagonist lmpL2. Dev Cell. 2015;33(1):36–46.

12. SongW, KirS, HongS, Hu Y, Wang X, Binari R, et al. Tumor-Derived Ligands Trigger TumorGrowth and Host Wasting via Differential MEKActivation. Dev Cell. 2019;48(2):277–86 e6.

13. Xu J, Liu Y, Yang F, Cao Y, Chen W, Li JSS, et al. Mechanistic characterization of a Drosophila model of paraneoplastic nephrotic syndrome. Nat Commun. 2024;15(1):1241.

14. Xu W, Li G, Chen Y, Ye X, Song W. A novel antidiuretic hormone governs tumour-induced renal dysfunction. Nature. 2023;624(7991):425–32.

15. Kwok SH, Liu Y, Bilder D, Kim J. Paraneoplastic renal dysfunction in fly cancer models driven by inflammatory activation of stem cells. Proc Natl Acad Sci USA. 2024;121 (42):e2405860121.

16. Cong F, Bao H, Wang X, Tang Y, Bao Y, Poulton JS, et al. Translocation of gut bacteria promotes tumor-associated mortality by inducing immune-activated renal damage. EMBO J. 2025;44(13):3586–613.

17. Yang SA, Portilla JM, Mihailovic S, Huang YC, Deng WM. Oncogenic Notch Triggers Neoplastic Tumorigenesis in a Transition-Zone-like Tissue Microenvironment. Dev Cell. 2019;49(3):461–72 e5.

18. De Gregorio E, Spellman PT, Tzou P, Rubin GM, Lemaitre B. The Toll and Imd pathways are the major regulators of the immune response in Drosophila. EMBO J. 2002;21(11):2568–79.

19. Iebba V, Totino V, Gagliardi A, Santangelo F, Cacciotti F, Trancassini M, et al. Eubiosis and dysbiosis: the two sides of the microbiota. New Microbiol. 2016;39(1):1–12.

20. Obata F, Fons CO, Gould AP. Early-life exposure to low-dose oxidants can increase longevity via microbiome remodelling in Drosophila. Nat Commun. 2018;9(1):975.

21. Kosakamoto H, Yamauchi T, Akuzawa-Tokita Y, Nishimura K, Soga T, Murakami T, et al. Local Necrotic Cells Trigger Systemic Immune Activation via Gut Microbiome Dysbiosis in Drosophila. Cell Rep. 2020;32(3):107938.

22. Kleino A, Silverman N. The Drosophila IMD pathway in the activation of the humoral immune response. Dev Comp Immunol. 2014;42(1):25–35.

23. Lee J, Song X, Hyun B, Jeon CO, Hyun S. Drosophila Gut Immune Pathway Suppresses Host Development-Promoting Effects of Acetic Acid Bacteria. Mol Cells. 2023;46(10):637–53.

24. Fast D, Duggal A, Foley E. Monoassociation with Lactobacillus plantarum Disrupts Intestinal Homeostasis in Adult Drosophila melanogaster. mBio. 2018;9(4).

25. Oi A, Shinoda N, Nagashima S, Miura M, Obata F. A nonsecretory antimicrobial peptide mediates inflammatory organ damage in Drosophila renal tubules. Cell Rep. 2025;44(1):115082.

26. Yamauchi T, Oi A, Kosakamoto H, Akuzawa-Tokita Y, Murakami T, Mori H, et al. Gut Bacterial Species Distinctively Impact Host Purine Metabolites during Aging in Drosophila. iScience. 2020;23(9):101477.

27. Akhgarjand C, Vahabi Z, Shab-Bidar S, Etesam F, Djafarian K. Effects of probiotic supplements on cognition, anxiety, and physical activity in subjects with mild and moderate Alzheimer’s disease: A randomized, double-blind, and placebo-controlled study. Front Aging Neurosci. 2022;14:1032494.

28. Hsu YC, Huang YY, Tsai SY, Kuo YW, Lin JH, Ho HH, et al. Efficacy of Probiotic Supplements on Brain-Derived Neurotrophic Factor, Inflammatory Biomarkers, Oxidative Stress and Cognitive Function in Patients with Alzheimer’s Dementia: A12-Week Randomized, Double-Blind Active-Controlled Study. Nutrients. 2023;16(1).

29. He X, Yan C, Zhao S, Zhao Y, Huang R, Li Y. The preventive effects of probiotic Akkermansia muciniphila on D-gaLactose/AlCl3 mediated Alzheimer’s disease-like rats. Exp Gerontol. 2022;170:111959.

30. Yang X, He X, Xu S, Zhang Y, Mo C, Lai Y, et al. Effect of Lacticaseibacillus paracasei strain Shirota supplementation on clinical responses and gut microbiome in Parkinson’s disease. Food Funct. 2023;14(15):6828–39.

31. Wang L, Zhao Z, Zhao L, Zhao Y, Yang G, Wang C, et al. Lactobacillus plantarum DP189 Reduces alpha-SYN Aggravation in MPTP-Induced Parkinson’s Disease Mice via Regulating Oxidative Damage, Inflammation, and Gut Microbiota Disorder. J Agric Food Chem. 2022;70(4):1163–73.

32. Pan S, Wei H, Yuan S, KongY, Yang H, Zhang Y, et al. Probiotic Pediococcus pentosaceus ameliorates MPTP-induced oxidative stress via regulating the gut microbiota-gut-brain axis. Front Cell Infect Microbiol. 2022;12:1022879.

33. Nurrahma BA, Tsao SP, Wu CH, Yeh TH, Hsieh PS, Panunggal B, Huang HY. Probiotic Supplementation Facilitates Recovery of 6-OHDA-lnduced Motor Deficit via Improving Mitochondrial Function and Energy Metabolism. Front Aging Neurosci. 2021;13:668775.

34. Kreuzer K, Reiter A, Birkl-Toglhofer AM, Dalkner N, Mork IS, Mairinger M, et al. The PROVIT Study-Effects of Multispecies Probiotic Add-on Treatment on Metabolomics in Major Depressive Disorder-A Randomized, Placebo-Controlled Trial. Metabolites. 2022;12(8).

35. Tian P, Chen Y, Zhu H, Wang L, Qian X, Zou R, et al. Bifidobacterium breve CCFM1025 attenuates major depression disorder via regulating gut microbiome and tryptophan metabolism: A randomized clinical trial. Brain Behav Immun. 2022;100:233–41.

36. Hashikawa-Hobara N, Otsuka A, Okujima C, Hashikawa N. Lactobacillus paragasseri OLL2809 Improves Depression-Like Behavior and Increases Beneficial Gut Microbes in Mice. Front Neurosci. 2022;16:918953.

37. Yu S, Luo F, Xu Y, Zhang Y, Jin LH. Drosophila Innate Immunity Involves Multiple Signaling Pathways and Coordinated Communication Between Different Tissues. Front Immunol. 2022;13:905370.

38. Chen Y, Xu W, Chen Y, Han A, Song J, Zhou X, Song W. Renal NF-kappaB activation impairs uric acid homeostasis to promote tumor-associated mortality independent of wasting. Immunity. 2022;55(9):1594–608 e6.

39. Torija MJ, Mateo E, Guillamon JM, Mas A. Identification and quantification of acetic acid bacteria in wine and vinegar byTaqMan-MGB probes. Food Microbiol. 2010;27(2):257–65.

40. Shi F, Liu Q, Yue D, Zhang Y, Wei X, Wang Y, Ma W. Exploring the effects of the dietary fiber compound mediated by a longevity dietary pattern on antioxidation, characteristic bacterial genera, and metabolites based on fecal metabolomics. Nutr Metab (Lond). 2024;21(1):18.

41. Koyle ML, Veloz M, Judd AM, Wong AC, Newell PD, Douglas AE, Chaston JM. Rearing the Fruit Fly Drosophila melanogaster Linder Axenic and Gnotobiotic Conditions. J Vis Exp. 2016(113).

42. StoreLli G, Defaye A, Erkosar B, Hols P, Royet J, Leulier F. Lactobacillus plantarum promotes Drosophila systemic growth by modulating hormonal signals through TOR-dependent nutrient sensing. Cell Metab. 2011;14(3):403–14.

